# The quagga mussel genome and the evolution of freshwater tolerance

**DOI:** 10.1101/505305

**Authors:** Andrew D. Calcino, André Luiz de Oliveira, Oleg Simakov, Thomas Schwaha, Elisabeth Zieger, Tim Wollesen, Andreas Wanninger

## Abstract

European freshwater dreissenid mussels evolved from marine ancestors during the Miocene approximately 30 million years ago and today include some of the most successful and destructive invasive invertebrate species of temperate freshwater environments. Here we sequenced the genome of the quagga mussel *Dreissena rostriformis* to identify evolutionary adaptations involved in embryonic osmoregulation. We found high gene expression levels of a novel subfamily of lophotrochozoan-specific aquaporin water channel, a vacuolar ATPase and a sodium/hydrogen exchanger during early cleavage, a period defined by the formation of inter-cellular fluid-filled ‘cleavage cavities’. Independent expansions of the lophotrochoaquaporin clade that coincide with at least five independent colonisation events of freshwater environments confirm their central role in freshwater adaptation. The pattern of repeated aquaporin expansion and the evolution of membrane-bound fluid-filled osmoregulatory structures in diverse taxa points to a fundamental principle guiding the evolution of freshwater tolerance that may provide a framework for future efforts towards invasive species control.

## MAIN TEXT

Molluscs evolved in the ocean, yet today there are approximately 5000 freshwater species worldwide originating from more than 40 independent colonisation events (1–3). While most freshwater mollusc species are gastropods (ca. 4000 species, 2), several bivalves (eg. *Dreissena rostriformis, Dreissena polymorpha, Limnoperna fortune* and *Corbicula fluminea*) have proven to be highly successful and ecologically disruptive ecosystem engineers of colonised environments (4–7).

Bivalves have invaded freshwater habitats on at least 11 occasions (3, 8). Despite the independence of these events, a feature common to freshwater species is a body fluid osmolarity that is actively maintained hyperosmotic to their environment (9, 10). This differs to marine and brackish water species which are commonly osmoconformers (11). Due to their large surface area to volume ratio, the issue of cellular osmoregulation is expected to be particularly acute for the eggs and embryos of broadcast spawners such as the quagga mussel *Dreissena rostriformis* (Fig. 1; Supplementary note SM 1). The evolution of mechanisms by the quagga mussel to withstand the harsh osmotic conditions of the freshwater environment during embryogenesis may be a contributing factor to their evolutionary success and their ability to rapidly spread through newly colonised habitats.

**Figure 1.**
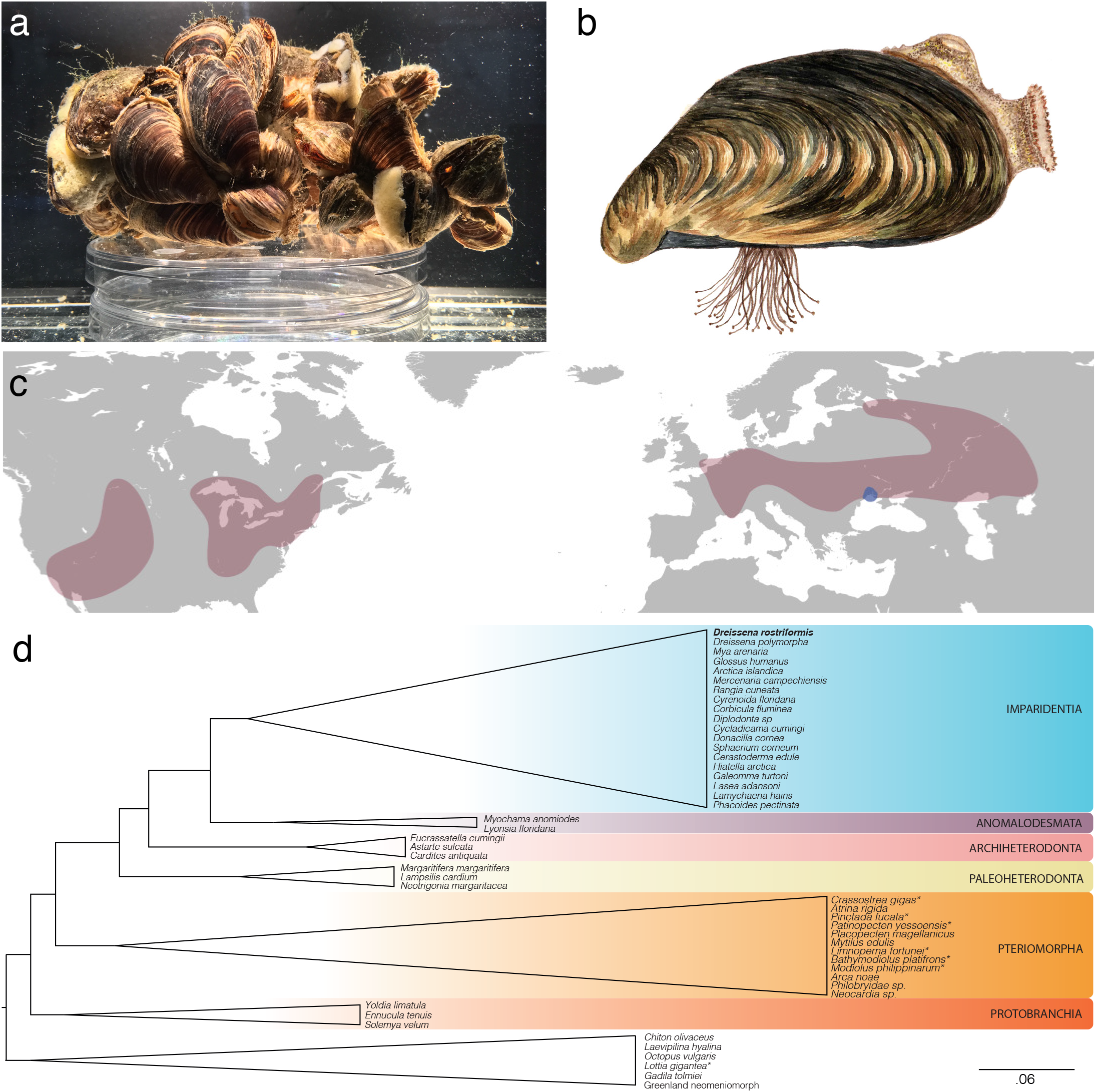
The quagga mussel, *Dreissena rostriformis*. **(a)** Quagga mussels form dense aggregations connected with strong byssal threads. Aggregates are often associated with other benthic species such as sponges. **(b)** Illustration of a single quagga mussel demonstrating the distinct banding pattern of the shell and the dense clump of byssus threads that enables them to adhere to both natural and manufactured substrates. **(c)** Global distribution of the quagga mussel highlighting native (blue) and colonised (red) habitats. **(d)** Condensed phylogeny of Bivalvia (58) based on a supermatrix composed of 47 molluscan taxa covering 1,377 orthogroups. The quagga mussel is positioned amongst the Imparidentia. Species with sequenced genomes are marked with a *. All nodes possess support values equal to 1.

Toidentify geneticsignatures that have underpinned the transition to freshwater environments, we sequenced, assembled and annotated the ~1.6 gigabase (Gb) genome of the quagga mussel *D. rostriformis* (Supplementary notes SM 1-2). High heterozygosity (~2.4%) and repeat content of the genome (Supplementary note SM 3) necessitated an assembly pipeline specifically designed for dealing with these issues, which are common to many wild-type non-inbred invertebrate species (Supplementary note SM 4; *12-15*). The resulting non-redundant haploid assembly covers 1.24 Gb with a scaffold N50 of 131.4 kilobases. The difference of ~360 megabases between the assembled genome and the predicted genome size is most likely explained by the collapsing or reduction of highly repetitive regions, as has been observed with other genome assemblies (16). Mapping of the paired end libraries back to the completed genome resulted in 94.5% realignment, confirming the integrity of the assembly.

Annotation of the assembly was performed through a combination of RNAseq-based transcriptome assemblies, homology data and *ab-inito* methods, resulting in the identification of 37,681 protein-coding gene models (Supplementary note SM 5). BUSCO analysis (v. 2.0, *17*) detected 928 (94.9%) of the 978 metazoan single copy orthologues and in total 32,708 (85.8%) of the gene models could be annotated based on homology to public databases. These results make the quagga mussel genome one of the most high quality and complete molluscan assemblies currently available and an important resource for metazoan genomics.

In order to maintain a body osmolarity above that of the surrounding medium, organisms require mechanisms wise diffuse across membranes, or to excrete excess water that passes into their cells. Likewise, the efflux of small solutes (eg. Na+, K+, Cl−) that would otherwise be lost to the environment must be prevented or balanced with uptake to avoid deteriorating osmotic conditions. An investigation of the genes encoding sodium, potassium, chloride and water transmembrane channels in *Dreissena* revealed an aquaporin, a vacuolar ATPase (v-ATPase) subunit a and a sodium/hydrogen exchanger (NHE) that are highly expressed in *Dreissena* eggs and early embryos (stages that lack distinct osmoregulatory organs) but not in those of the marine oyster *Crassostrea gigas* (Fig. 2a; Supplementary note SM6).

**Figure 2.**
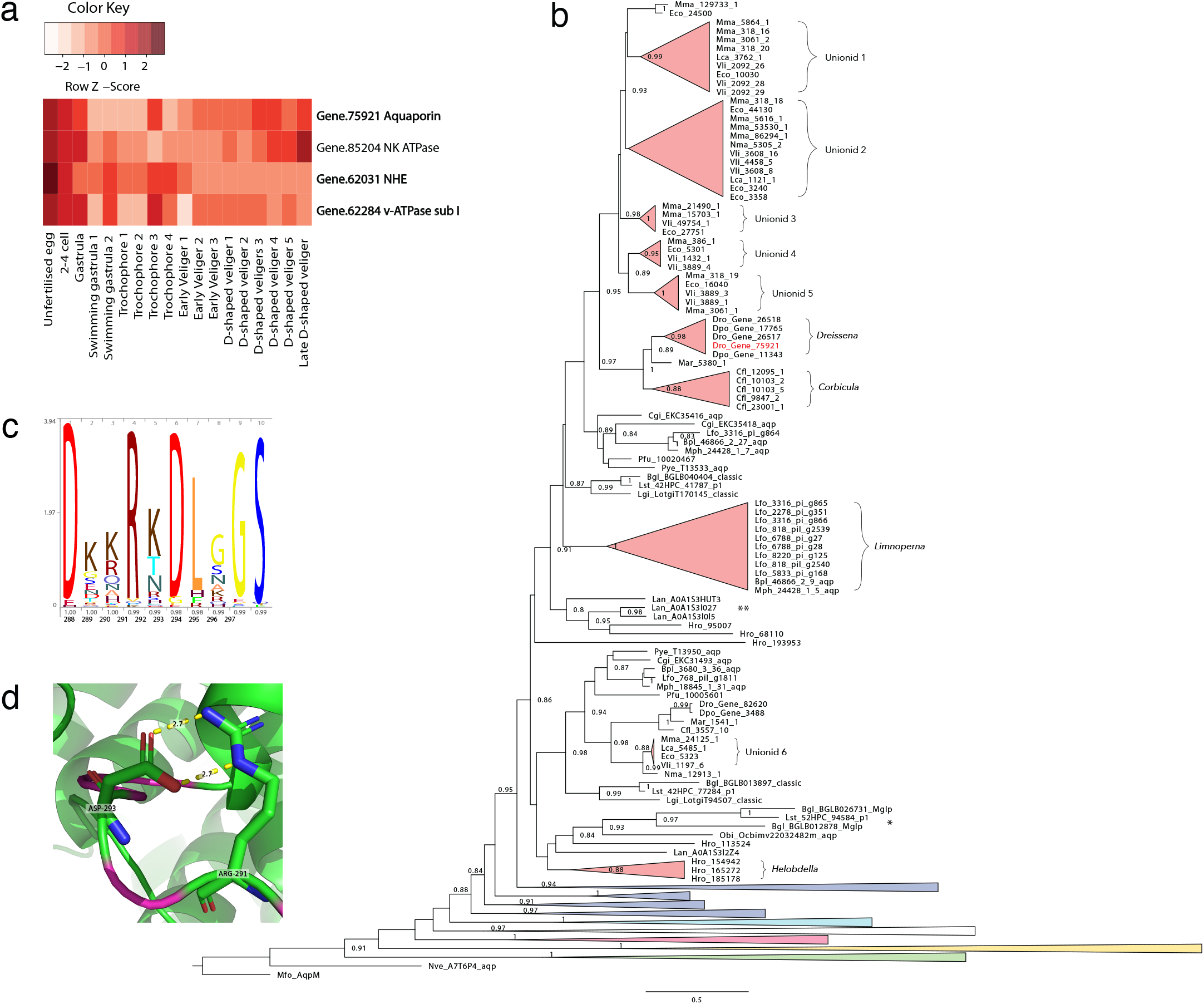
Embryonic osmotic regulatory genes. **(a)** Heat map of expression of candidate osmoregulatory genes highly expressed during embryogenesis prior to the free-swimming stage. Three genes (highlighted in bold) are highly expressed in *Dreissena* but not in similar stages of the marine oyster *Crassostrea gigas* (see SM 6.2). **(b)** Phylogenetic tree of aquaporins with emphasis on classical lophotrochoaquaporins (red) with other classes (green - aquaglycer-oporins, yellow - unorthodox, pink - EGLPs, white - undescribed annelid/brachiopod clade, light blue - aquaamoni-aporins, blue - classical aquaporins) collapsed. Note the independent expansions associated with the freshwater lineages *Dreissena, Corbicula, Limnoperna*, the unionid mussels and the annelid leech *Helobdella*. * indicates a clade of long branch freshwater gastropod sequences previously annotated as malacoglyceroporins. ** indicates an expanded clade of marine brachiopod sequences. Support values under 0.8 are not shown. **(c)** Peptide logo of the highly charged lophotrochoaquaporin loop D with occupancy and amino acid position indicated on the x-axis respectively. **(d)** Predicted structure of the *Dreissena* lophotrochoaquaporin *Dro.75921* loop D (magenta) wrapped to *Bos taurus AQP1* (PDB: 1j4n.1) showing the predicted salt bridge formed between Arg-291 and Asp-293.

Aquaporins are a class of highly selective transmembrane passive water transporters that in animals fall into one of four clades (classical, aquaamoniaporins, aquaglyceroporins or unorthodox/S-aquaporins) based on their function and their capacity to transport specific solutes in addition to or instead of water (18). A fifth group specific to insects, the entomoglyceroporins (EGLPs), have been previously assigned to the classical aquaporins (18), however our results indicate they may represent a sister group. In humans, aquaporins are associated with a number of pathologies and are important components of the eye, blood-brain-barrier and kidney (19). In molluscs, the diversity and function of aquaporins are less well described although mollusc-specific aquaporin clades have been identified (20, 21).

A phylogeny of 291 aquaporin genes including 168 bivalve sequences from 15 species spanning both marine and freshwater taxa demonstrates a distinctive pattern of independent aquaporin gene expansion in freshwater bivalve species (Fig. 2b; Supplementary note SM 7). A previously unidentified class of lophotrochozoan-specific classical aquaporins (hereafter referred to as lophotrochoaquaporins) that includes the highly expressed *Dreissena* genes appears to have expanded on at least four occasions, coinciding with the freshwater bivalves *Dreissena, Corbicula fluminea, Limnoperna fortunei* and the Unionidae (a species-rich monophylectic taxon of freshwater paleoheterodont bivalves). In contrast, none of the marine bivalves show evidence of such an expansion. *Dreissena, Corbicula* and *Limnoperna* represent independent invasions of freshwater habitats and all three have close marine relatives (Supplementary note SM 1; *8, 15*). The lineage-specific aquaporin duplications in these three species reflect this evolutionary history while the aquaporin expansion in the paleoheterodonts appears to have occurred after the divergence of the freshwater unionids from their marine ancestors, but before unionid speciation (Fig. 2b). Outside of the bivalves, the freshwater annelid leech *Helobdella robusta* also has an expanded set of lophotrochoaquaporins while the freshwater gastropods *Limnaea stagnalis* and *Biomphalaria glabrata* possess long branch lengths without evidence of expansions (Fig. 2b).

In addition to facilitating water transport across osmotic gradients, hydrostatic gradients may also influence the directional transport of water by aquaporin channels. In the absence of an osmotic pressure gradient, rat *AQP1* channels mediate transmembrane water transport across a hydrostatic pressure gradient (22) while human *AQP1* expressed in *Xenopus* oocytes exhibits reversible gating dependent on the surface tension of the membrane (23). In plants, membrane tension-dependent gating of the grapevine aquaporin TIP2;1 has also been observed under hypotonic conditions (24).

Modelling of the three-dimensional conformation of the embryonically expressed quagga mussel lophotocho-aquaporin orthologue *Dro.75921* with SWISS-MODEL (25) revealed high structural conservation of the transmembrane regions to other classical aquaporins and structural variability predominantly confined to loops A, C and E (Supplementary note SM 8). The exception to this is *AQP1*, which also shows evidence of structural similarity with *Dro.75921* through loop D. In lophotrochoaquaporins, loop D is highly charged and appears to be highly conserved with two of the most conserved amino acids (Arg-291 and Asp-293 in *Dro.75921*) also predicted to form a salt bridge (Fig. 2d; 26). In addition to the conductance of water through individual subunit pores, *AQP1* conducts Na+, K+ and Cs+ cations through the central tetrameric pore which is gated by the highly charged cytoplasmic loop D (27–29). Increasing evidence that water and ion conductance of aquaporins can be influenced by factors such as pH (30–33), membrane tension (23) and hydrostatic pressure (22) suggests that a complex set of dynamic, localised parameters may interact to influence aquaporin activity.

A conspicuous feature of the early cleavage stages of some freshwater and terrestrial molluscs is the formation of a large lens-shaped, fluid-filled cavity between dividing cells (34, 35). Following cell division, small cavities begin to appear along the cell-cell border, gradually coalescing until a single large cavity remains. The contents of the cleavage cavity are then rapidly discharged to the environment before a new cavity begins to form (Fig. 3, a-d; Supplementary note SM 8). This process is typically repeated two to four times for each of the first three cell divisions. At the 8-cell stage, intercellular cleavage cavities become less apparent and a single central blastocoel forms (Supplementary note SM 9). This, too, undergoes periods of gradual inflation and rapid discharge up until at least the swimming gastrula stage.

**Figure 3.**
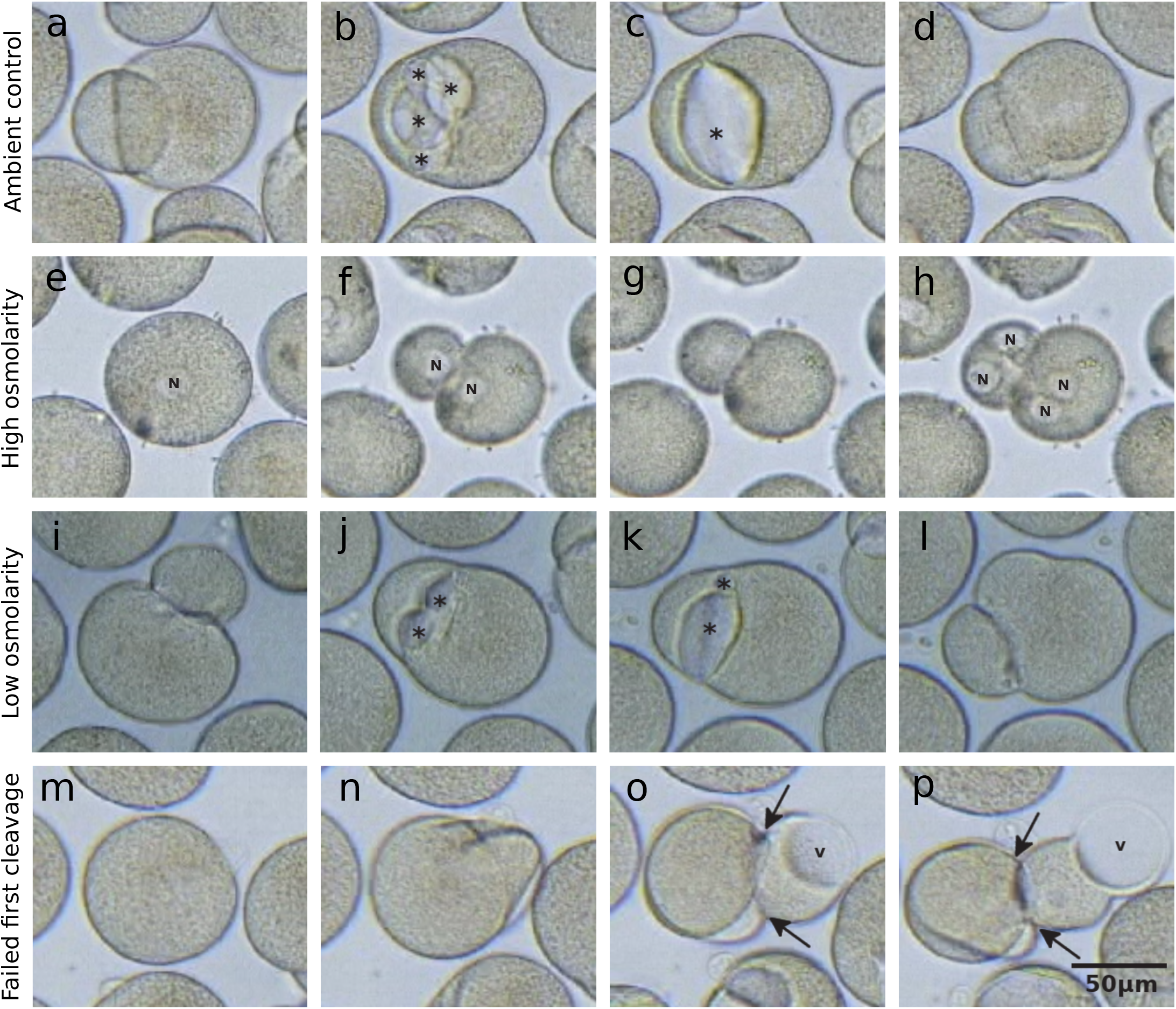
Cleavage cavities in developing *Dreissena* embryos. **(a-d)** Formation of cleavage cavities during the first embryonic cleavage under ambient conditions. **(e-h)** First and second embryonic cleavages under high salinity demonstrating the lack of cleavage cavity formation. **(i-l)** Formation of cleavage cavities during the first embyronic cleavage under low salinity, **(m-n)** Failed first cleavage leading to rupture of the fertilisation envelope and the extrusion of an amoeboid projection with a large vacuole. * indicates cleavage cavities, n indicates nuclei, v indicates intracellular vacuoles and arrows indicate fertilisation envelope ruptures.

Small cleavage cavities are also present in marine molluscs, however, these remain small and fail to coalesce (36). In contrast, in freshwater and terrestrial molluscs, the formation of a wide cleavage cavity has been observed in multiple phylogenetically disparate species (34–39). Outside of the molluscs, the freshwater annelid leech *Helobdella* also forms cleavage cavities which gradually inflate, coalesce and rapidly discharge in the same manner as those of molluscs (40, 41). The freshwater bryozoan *Paludicella articulata* may also form a structure reminiscent of molluscan cleavage cavities (42). In vertebrates, rat hepatocyte couplets induced to secrete bile through the application of the choleretic dibutyryl cAMP rapidly shunt intracellular *AQP8* to the membranes at the cell-cell interface (43). This induces the flow of water in the presence or absence of an osmotic gradient, forming a fluid-filled intercellular structure (the canaliculus) with striking resemblance to *Dreissena* cleavage cavities.

To confirm a role for the *Dreissena* cleavage cavity formation in osmotic regulation, developing embryos were subjected to hyperosmotic conditions (Supplementary note SM 9). In the majority of circumstances this resulted in a complete absence of cleavage cavity formation, while in a few individuals, cleavage cavity formation remained but was greatly reduced in size compared to those grown in ambient conditions (Fig. 3, e-h). Under reduced salinity, cleavage cavities formed normally (Fig. 3, i-l). This is consistent with results from other cleavage cavity forming species (35, 37, 44, 45).

Like embryonic molluscs, freshwater protozoans require mechanisms to cope with the osmotic influx of water across their membranes in the absence of complex multicellular excretory organs. Similar to lophotrochozoan cleavage cavities, protozoan contractile vacuoles are membrane-bound structures that gradually fill with excess cellular water before rapidly discharging it to the exterior (46, 47). Additionally, both cleavage cavities and contractile vacuoles are associated with aquaporins and vacuolar ATPases (Fig. 2; Supplementary note SM 6; 48). Our data provide the first evidence for the central role of aquaporins and vacuolar ATPases in a lophotrochozoan osmoregulatory process which is similar to that of protozoan contractile vacuoles. The repeated aquaporin expansions in freshwater lophotrochozoans and the importance of aquaporins and v-ATPases to protozoan osmoregulatory structures suggests that this molecular machinery may have been co-opted by both the lophotrochozoan cleavage cavity and the protozoan contractile vacuole excretory systems.

The NHE that is highly expressed in *Dreissena* embryogenesis is a member of the plasma-membrane NHEs and thus is likely required for the recovery of sodium lost to the environment across the plasma membrane. Neither the v-ATPases nor the PM-NHEs have undergone an evolutionary pattern of expansion akin to that of the aquaporins in which repeated colonisation events of freshwater environments are associated with gene family expansions (Fig. 2; Supplementary note SM 7). This further highlights a likely central role for aquaporins in lophotrochozoan osmoregulation.

The co-occurrence of cleavage cavity formation with high aquaporin and v-ATPase subunit a expression levels indicates a role for these proteins in osmoregulation during early *Dreissena* embryogenesis similar to that in protozoan contractile vacuoles. While v-ATPases are most well-known for the acidification of intracellular vesicles, their numerous roles in the acidification of extracellular spaces are now well established (49, 50). It is also significant that the only v-ATPase subunit that is highly upregulated during cleavage cavity formation is an orthologue of subunit a as it is this subunit that is responsible for subcellular targeting of the entire v-ATPase complex (51–53). The protonation of the cleavage cavity that is predicted to result from v-ATPase activity may influence aquaporin-mediated water or ion conductance in a pH-dependent manner as has been observed for several mammalian aquaporins (30–33).

In protozoans, the mechanism of contractile vacuole contraction remains elusive. Contractile elements such as actin-myosin-based systems do not appear to be associated with contractile vacuoles, leading some authors to suggest that “contractile” vacuoles are not actually contractile (46). A more recent hypothesis posits that tension built up in the membrane of the contractile vacuole may provide the force required for fluid discharge (54).

In many animals, post-fertilisation physicochemical modifications that transform the vitelline envelope into the fertilisation envelope typically result in increased rigidity and stiffness (55–57). While such modifications have not been well described in molluscs, we observed that *Dreissena* embryos that fail to cleave or to form cleavage cavities under ambient conditions appear to be at increased risk of fertilisation envelope rupture (Fig. 3, m-p). In contrast, unfertilised *Dreissena* eggs gradually increase in volume, presumably due to the osmotic influx of water, without such a marked increase in susceptibility to vitelline envelope rupture (Supplementary note SM 9).

We suggest a mechanism for *Dreissena* cleavage cavity filling and discharge whereby the mechanical properties of the fertilisation envelope restricts cell expansion in a functionally analogous manner to that of the cell wall of plants. Under such circumstances, an increased turgor pressure induced by the osmotic influx of water triggers the flow of water through the aquaporin channels down a hydrostatic pressure gradient into the developing cleavage cavity. The rapid expulsion of the contents of the cleavage cavity is thus mediated by tension stored in the fertilisation envelope in a manner similar to the tension stored in the membranes of protozoan contractile vacuoles (54).

The deployment of lophotrochoaquaporin water transport channels during early embryogenesis appears to have enabled *Dreissena* to regulate intercellular water levels as early as the two-cell stage of development. The ability of *Dreissena* embryos to excrete excess water via the formation of cleavage cavities is a function that in adult animals is usually reserved for complex organs such as nephridia or kidneys. It is likely that the evolution of this embryonic osmotic regulatory mechanism involving aquaporin-mediated cleavage cavity formation was a crucial step in the adaptation of the quagga mussel - and likely many other aquatic animals - to freshwater environments.

## ETHICS APPROVAL AND CONSENT TO PARTICIPATE

No specific ethics approval was required for this project.

## AUTHORS’ CONTRIBUTIONS

ADC contributed to the design of the project, field work, genome construction and annotation, analysis and interpretation of the data, laboratory procedures and manuscript preparation. ALDO contributed to the phylogenetic and phylogenomic analyses, OS contributed to data analysis and drafting of the manuscript, TS contributed to microscopy and image analysis, TW contributed to field and lab work and AW contributed to the design of the project, analysis and interpretation of the data and manusript preparation.

## ACKNOWLEDGEMENTS

We would like to thank Paulina Tapia for her quagga mussel water colour artwork and photography in Figure 1, and for the design of the document.

## REFERENCES

1. E. V Balian, H. Segers, C. Lévèque, K. Martens, The freshwater animal diversity assessment: an overview of the results. Hydrobiologia. 595, 627–637 (2008).

2. E. E. Strong, O. Gargominy, W. F. Ponder, P. Bouchet, Global diversity of gastropods (Gastropoda; Mollusca) in freshwater. Hydrobiologia. 595, 149–166 (2008).

3. D. L. Graf, Patterns of freshwater bivalve global diversity and the state of phylogenetic studies on the Unionoida, Sphaeriidae, and Cyrenidae. Am. Malacol. Bull. 31, 135–153 (2013).

4. S. Werner, K.-O. Rothhaupt, Effects of the invasive bivalve *Corbicula fluminea* on settling juveniles and other benthic taxa. J. North Am. Benthol. Soc. 26, 673–680 (2007).

5. R. Sousa, J. L. Gutiérrez, D. C. Aldridge, Non-indigenous invasive bivalves as ecosystem engineers. Biol. Invasions. 11, 2367–2385 (2009).

6. G. Darrigran, C. Damborenea, Ecosystem engineering impact of *Limnoperna fortunei* in South America. Zoolog. Sci. 28, 1–7 (2011).

7. K. M. DeVanna, P. M. Armenio, C. A. Barrett, C. M. Mayer, Invasive ecosystem engineers on soft sediment change the habitat preferences of native mayflies and their availability to predators. Freshw. Biol. 56, 2448–2458 (2011).

8. D. J. Combosch et al., A family-level tree of life for bivalves based on a Sanger-sequencing approach. Mol. Phylogenet. Evol. 107, 191–208 (2017).

9. L. E. Deaton, Ion regulation in freshwater and brackish water bivalve mollusks. Physiol. Zool. 54, 109–121 (1981).

10. T. H. Dietz, et al., Osmotic and ionic regulation of North American zebra mussels (*Dreissena polymorpha*). Am. Zool. 36, 364–372 (1996).

11. M. B. Griffith, Toxicological perspective on the osmoregulation and ionoregulation physiology of major ions by freshwater animals: teleost fish, Crustacea, aquatic insects, and Mollusca. Environ. Toxicol. Chem. 36, 576–600 (2017).

12. Q. Xia et al., A draft sequence for the genome of the domesticated silkworm (*Bombyx mori*). Science. 306, 1937–1940 (2004).

13. G. Zhang et al., The oyster genome reveals stress adaptation and complexity of shell formation. Nature. 490, 49–54 (2012).

14. Q. Cong, D. Borek, Z. Otwinowski, N. V Grishin, Tiger swallowtail genome reveals mechanisms for speciation and caterpillar chemical defense. Cell Rep. 10, 910–919 (2015).

15. M. Uliano-Silva et al., A hybrid-hierarchical genome assembly strategy to sequence the invasive golden mussel *Limnoperna fortunei*. Gigascience. 7, gix128 (2018).

16. S. Wang et al., Scallop genome provides insights into evolution of bilaterian karyotype and development. Nat. Ecol. Evol. 1, 0120 (2017).

17. A. Simão, R. M. Waterhouse, P. Ioannidis, E. V Kriventseva, E. M. Zdobnov, BUSCO: assessing genome assembly and annotation completeness with single-copy orthologs. Bioinformatics. 31, 3210–3212 (2015).

18. R. N. Finn, J. Cerdà, Evolution and functional diversity of aquaporins. Biol. Bull. 229, 6–23 (2015).

19. A. S. Verkman, Aquaporins at a glance. J. Cell Sci. 124, 2107–2112 (2011).

20. E. Kosicka, D. Grobys, H. Kmita, A. Lesicki, J. R. Pieńkowska, Putative new groups of invertebrate water channels based on the snail *Helix pomatia L*. (Helicidae) MIP protein identification and phylogenetic analysis. Eur. J. Cell Biol. 95, 543–551 (2016).

21. D. J. Colgan, R. P. Santos, A phylogenetic classification of gastropod aquaporins. Mar. Genomics. 38, 59–65 (2018).

22. T. Nguyen et al., Aquaporin-1 facilitates pressure-driven water flow across the aortic endothelium. Am. J. Physiol. Heart Circ. Physiol. 308, H1051–64 (2015).

23. M. Ozu, R. A. Dorr, F. Gutiérrez, M. Teresa Politi, R. Toriano, Human *AQP1* is a constitutively open channel that closes by a membrane-tension-mediated mechanism. Biophys. J. 104, 85–95 (2013).

24. L. Leitão, C. Prista, M. C. Loureiro-Dias, T. F. Moura, G. Soveral, The grapevine tonoplast aquaporin TIP2;1 is a pressure gated water channel. Biochem. Biophys. Res. Commun. 450, 289–294 (2014).

25. A. Waterhouse et al., SWISS-MODEL: Homology modelling of protein structures and complexes. Nucleic Acids Res. 46, W296–W303 (2018).

26. S. Costantini, G. Colonna, A. M. Facchiano, ESBRI: A web server for evaluating salt bridges in proteins. Bioinformation. 3, 137–138 (2008).

27. T. L. Anthony et al., Cloned human aquaporin-1 is a cyclic GMP-gated ion channel. Mol. Pharmacol. 57, 576–88 (2000).

28. J. Yu, A. J. Yool, K. Schulten, E. Tajkhorshid, Mechanism of gating and ion conductivity of a possible tetrameric pore in aquaporin-1. Structure. 14, 1411–1423 (2006).

29. M. Kourghi, M. L. De Ieso, S. Nourmohammadi, J. V. Pei, A. J. Yool, Identification of loop D domain amino acids in the human aquaporin-1 channel involved in activation of the ionic conductance and inhibition by AqB011. Front. Chem. 6, 1–12 (2018).

30. M. Yasul et al., Rapid gating and anion permeability of an intracellular aquaporin. Nature. 402, 184–187 (1999).

31. D. Alberga et al., A new gating site in human aquaporin-4: Insights from molecular dynamics simulations. Biochim. Biophys. Acta, Biomembr. 1838, 3052–3060 (2014).

32. S. Kaptan et al., H95 Is a pH-dependent gate in aquaporin 4. Structure. 23, 2309–2318 (2015).

33. C. Rodrigues et al., Rat aquaporin-5 is pH-gated induced by phosphorylation and is implicated in oxidative stress. Int. J. Mol. Sci. 17 (2016)

34. J. Meisenheimer, Entwicklungsgeschichte von Dreissensia polymorpha Pall. (1900).

35. C. A. Kofoid, On the early development of *Limax*. Bull. Museum Comp. Zool. 27, 33–118 (1895).

36. C. Raven, Morphogenesis: The analysis of molluscan development (Pergamon, ed 2, 1966), pp. 64–109.

37. C. Raven, The development of the egg of *Limnea stagnalis L*. from the first cleavage till the trochophore stage, with special reference to its “chemical embryology.” Arch. Néerl. Zool. 7, 353–434 (1936).

38. T. Kawano et al., Observation of some key stages of the embryonic development of *Biomphalaria straminea* (Dunker, 1848) (molluska, planorbidae). Invertebr. Reprod. Dev. 46, 85–91 (2004).

39. H. H. Taylor, The ionic and water relations of embryos of *Lymnaea stagnalis*, a freshwater pulmonate mollusc. J. Exp. Biol. 69, 143–172 (1977).

40. D. C. Lyons, D. A. Weisblat, D quadrant specification in the leech *Helobdella*: Actomyosin contractility controls the unequal cleavage of the CD blastomere. Dev. Biol. 334, 46–58 (2009).

41. D. A. Weisblat, D. H. Kuo, Developmental biology of the leech *Helobdella*. Int. J. Dev. Biol. 58, 429–443 (2014).

42. S. O. Martha, Y. Afshar, A. N. Ostrovsky, T. Schwaha, T. S. Wood, “ Variation of the tentacles in Paludicella: the unfinished work of the German bryozoologist and embryologist Fritz Braem” in Annals of Bryozoology 6: aspects of the history of research on bryozoans. (International Bryozoology Association, 2018), p 45–74.

43. R. C. Huebert, P. L. Splinter, F. Garcia, R. A. Marinelli, N. F. Larusso, Expression and localization of aquaporin water channels in rat hepatocytes. J. Biol. Chem. 277, 22710–22717 (2002).

44. C. Raven, H. Klomp, The osmotic properties of the egg of *Limnaea stagnalis L*. Proc. Kon. Ned. Akad. Wet. Amsterd. 49, 101–109 (1946).

45. H. H. Taylor, The ionic properties of the capsular fluid bathing embryos of *Lymnaea stagnalis* and *Biomphalaria sudanica* (Mollusca: Pulmonata). J. Exp. Biol. 59, 543–564 (1973).

46. R. D. Allen, The contractile vacuole and its membrane dynamics. BioEssays. 22, 1035–1042 (2000).

47. H. Plattner, The contractile vacuole complex of protists-New cues to function and biogenesis. Crit. Rev. Microbiol. 41, 218–227 (2015).

48. E. Nishihara et al., Presence of aquaporin and V-ATPase on the contractile vacuole of *Amoeba proteus*. Biol. Cell. 100, 179–188 (2008).

49. S. Breton, D. Brown, New insights into the regulation of V-ATPase-dependent proton secretion. AJP Ren. Physiol. 292, F1–F10 (2006).

50. S. Pamarthy, A. Kulshrestha, G. K. Katara, K. D. Beaman, The curious case of vacuolar ATPase: Regulation of signaling pathways. Mol. Cancer. 17, 1–9 (2018).

51. S. Kawasaki-Nishi, K. Bowers, T. Nishi, M. Forgac, T. H. Stevens, The amino-terminal domain of the vacuolar proton-translocating ATPase a subunit controls targeting and in vivo dissociation, and the carboxyl-terminal domain affects coupling of proton transport and ATP hydrolysis. J. Biol. Chem. 276, 47411–47420 (2001).

52. M. Forgac, Vacuolar ATPases: Rotary proton pumps in physiology and pathophysiology. Nat. Rev. Mol. Cell Biol. 8, 917–929 (2007).

53. M. Toei, R. Saum, M. Forgac, Regulation and isoform function of the V-ATPases. Biochemistry. 49, 4715–4723 (2010).

54. T. Tani, T. Tominaga, R. D. Allen, Y. Naitoh, Development of periodic tension in the contractile vacuole complex membrane of *Paramecium* governs its membrane dynamics. Cell Biol. Int. 26, 853–860 (2002).

55. Y. Hiramoto, Mechanical properties of the surface of the sea urchin egg at fertilization and during cleavage. Exp. Cell Res. 89, 320–326 (1974).

56. G. L. Gerton, J. L. Hedrick, The vitelline envelope to fertilization envelope conversion in eggs of *Xenopus laevis*. Dev. Biol. 116, 1–7 (1986).

57. M. Khalilian, M. Navidbakhsh, M. R. Valojerdi, M. Chizari, E. Yazdi, Estimating Young’ s modulus of zona pellucida by micropipette aspiration in combination with theoretical models of ovum. J. R. Soc. Interface. 7, 687–694 (2010).

58. V. L. Gonzalez et al., A phylogenetic backbone for Bivalvia: an RNA-seq approach. Proc. R. Soc. B Biol. Sci. 282, 20142332–20142332 (2015).

