## Supplementary Material for "The quagga mussel genome and the evolution of freshwater tolerance"

### Materials and Methods

#### SM 1 - Taxonomy and phylogenetics of the quagga mussel

This history of the taxonomic classification of the genus *Dreissena* is long, complex and incomplete. Since its description in the nineteenth century, the quagga mussel has been synonymised as *Dreissena rostriformis*, *Dreissena bugensis* and *Dreissena rostriformis bugensis* (*1*, *2*). Reported differences between *D. rostriformis* and *D. bugensis* include the depth at which they are found and the salinity of their native habitats. However, attempts to discriminate the two on the basis of morphology have proven difficult due to the high level of intraspecific relative to interspecific variation. More recent efforts to discriminate the two on the basis of molecular markers (*COI*, *16S* rRNA) have concluded that rather than a distinct species, *D. bugensis* is likely to be a subspecies of *D. rostriformis* (*2*). This has led to the reclassification of the shallow freshwater form as *Dreissena rostriformis bugensis* due to taxonomic conventions stipulating the first described species name (*Dreissena rostriformis*, ANDRUSOV 1839) be retained over the later described species (*Dreissena bugensis*, ANDRUSOV 1867).

The World Register of Marine Species (WORMS) database currently recognises both *D. rostriformis* and *D. bugensis* as accepted species names while the status of *D. rostriformis bugensis* is ‘unaccepted’. In Austria, where the specimens for this project were collected, no record of the existence of the quagga mussel has yet been reported in the literature, however quagga mussels have been identified in the Danube river in Romania (*3*) and Serbia (*4*) and in the river Main in Germany which is connected to the Danube through the Main-Danube canal (*5*). To test the taxonomic status of the *D. rostriformis* sampled here, we performed a phylogeny based on *COI* sequences using all dreissenids available from Barcode of Life Database (BoLD), and those used by Therriaut et al. (2004) to discern the two clades (Fig. S1).

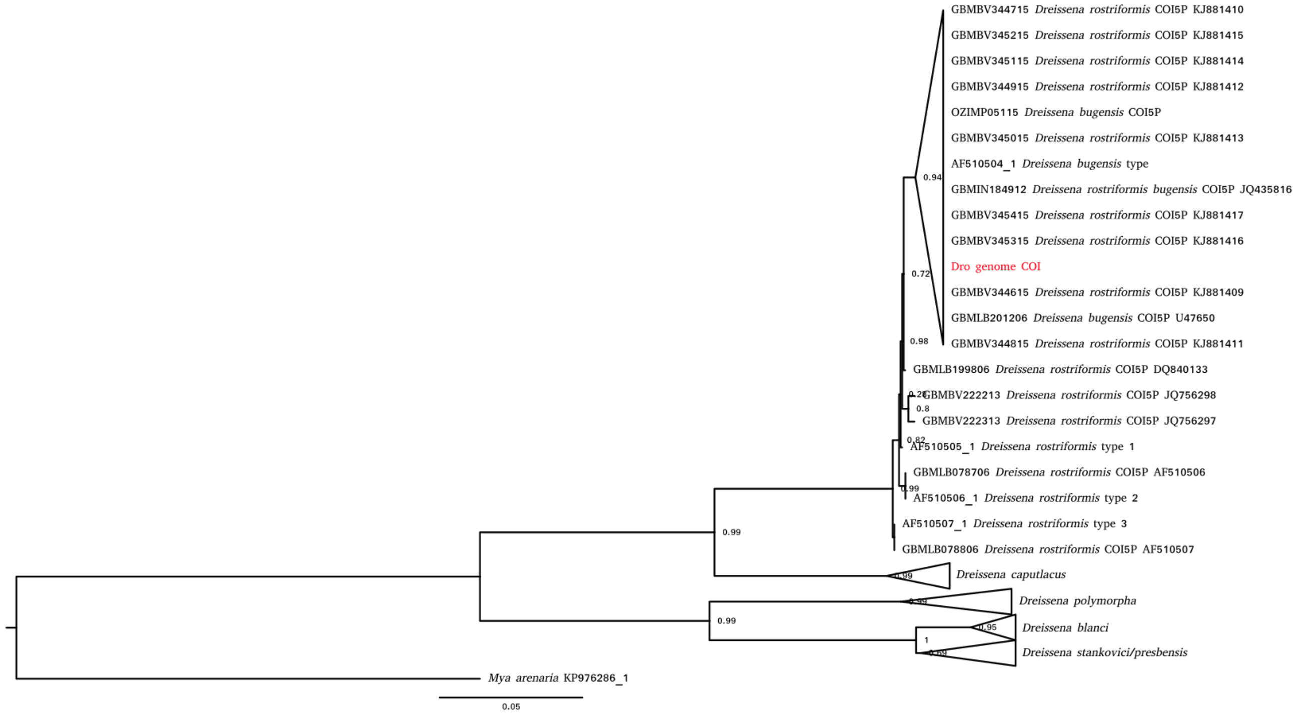

**Figure S1** - Dreissenid *COI* phylogeny. The sequence in red is the *COI* from the genome sequenced here, sequences with 'type' in the name were obtained by Therriault et al. (2004) and the remaining sequences were obtained from the BoLD database. *D. stankovici* and *D. presbensis* are likely to represent a single species called *Dreissena carinata* (Dunker, 1853).

Our results support the discontinuation of *D. bugensis* as a species distinct from *D. rostriformis* and they also indicate that the species names allocated to the quagga mussel samples in the BoLD database (*D. rostriformis, D. bugensis, D. rostriformis bugensis*) do not represent distinct genetic clades. As such, the preferred name is *D. rostriformis* as it this the oldest of the three names. A single well supported clade within the *D. rostriformis* branch that includes the sample sequenced here in addition to the *D. bugensis* sample collected by Therriault (2004) was identified, suggesting that the shallow freshwater form may represent a genetically distinct group, although more dedicated sampling will be required to confirm this. This analysis was unable to resolve the distinction between the BoLD *D. presbensis* and *D. stankovici COI* sequences. Neither of these species names are marked as 'accepted' on the WORMS database and it is likely that both are synonyms for *Dreissena carinata* (Dunker, 1853).

In examining the *16S* rRNA sequenced of *D. rostriformis* and *D. bugensis*, Therriault et al. (2004) identified a single nucleotide difference between the two forms which could be used as a diagnostic identification tool by cleaving PCR products with the restriction enzymes Msp I or HpaII. The *16S* sequence from the sample sequenced here is consistent with *D. rostriformis* however, as with the COI analysis, more dedicated sampling will be required to confirm this as a diagnostic feature (Fig. S2).

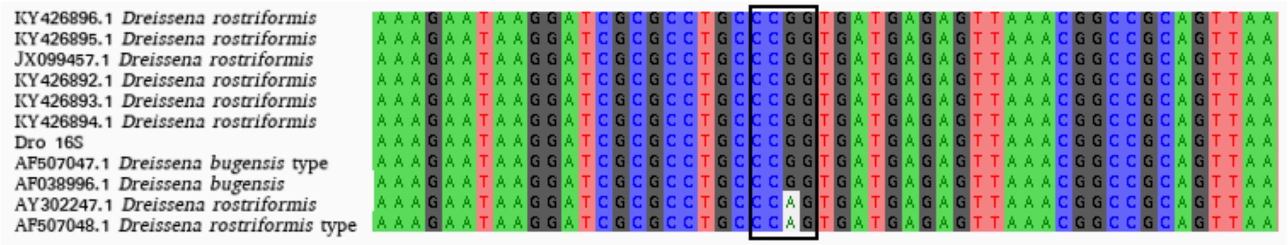

**Figure S2 -** Multiple sequence alignment of *16S* rRNA. The *16S* rRNA from the genome sequenced here is named *Dro 16S*, sequences with 'type' in the name were obtained by Therriault et al. (2004) and the remaining sequences were downloaded from NCBI. The box highlights a motif identified by Therriault et al. (2004) as diagnostic for discerning *bugensis* (CCGG) from other *D. rostriformis* clades (CCAG).

### SM 2 - Phylogenomics of the quagga mussel

To confirm the phylogenetic position of the quagga mussel and the other species used for comparative analyses in this study, a phylogenomic tree was produced (Fig. S3). A total of 1,377 curated orthogroups obtained from 40 molluscan taxa, including 34 bivalves (*6*), were downloaded and used to build profile-hidden Markov models (pHMMs) and multiple sequence alignments (MSAs) to extend the orthologue groups using HaMStr (*7*). Multiple sequence alignments were generated with mafft (*8*) and pHMMs built with hmmbuild from the HMMER3 package (*9*). Protein-coding sequences from five publicly available genomes (*Bathymodiolus platifrons*, *Crassostrea gigas*, *Limnoperna fortunei*, *Modiolus philippinarum*, and *Patinopecten yessoensis*), one transcriptome (*Dreissena polymorpha*, see SM 7 for assembly details), and from the quagga mussel (see SM 5) were searched against the 1,377 pHMMs in HaMStR using the default options with the -representative option. Each potential candidate orthologue was then rechecked with reciprocal BLAST against the reference taxa *Lottia gigantea*, *Corbicula fluminea*, *Ennucula tenuis, Solemya velum*, *Mytilus edulis*, *Mya arenaria*, and discarded if it the reciprocal hit was not fulfilled. The extended 1377 orthogroups were concatenated into a super matrix with FASconCAT v1.11 (*10*), and the phylogeny inferred with FastTree (*11*) using default parameters and -lg model of amino acid substitution. We find that the closest relative to *Dreissena* is the soft shell clam *Mya arenaria* and that the *Dreissena* lineage has the longest branch length of the Imparidentia.

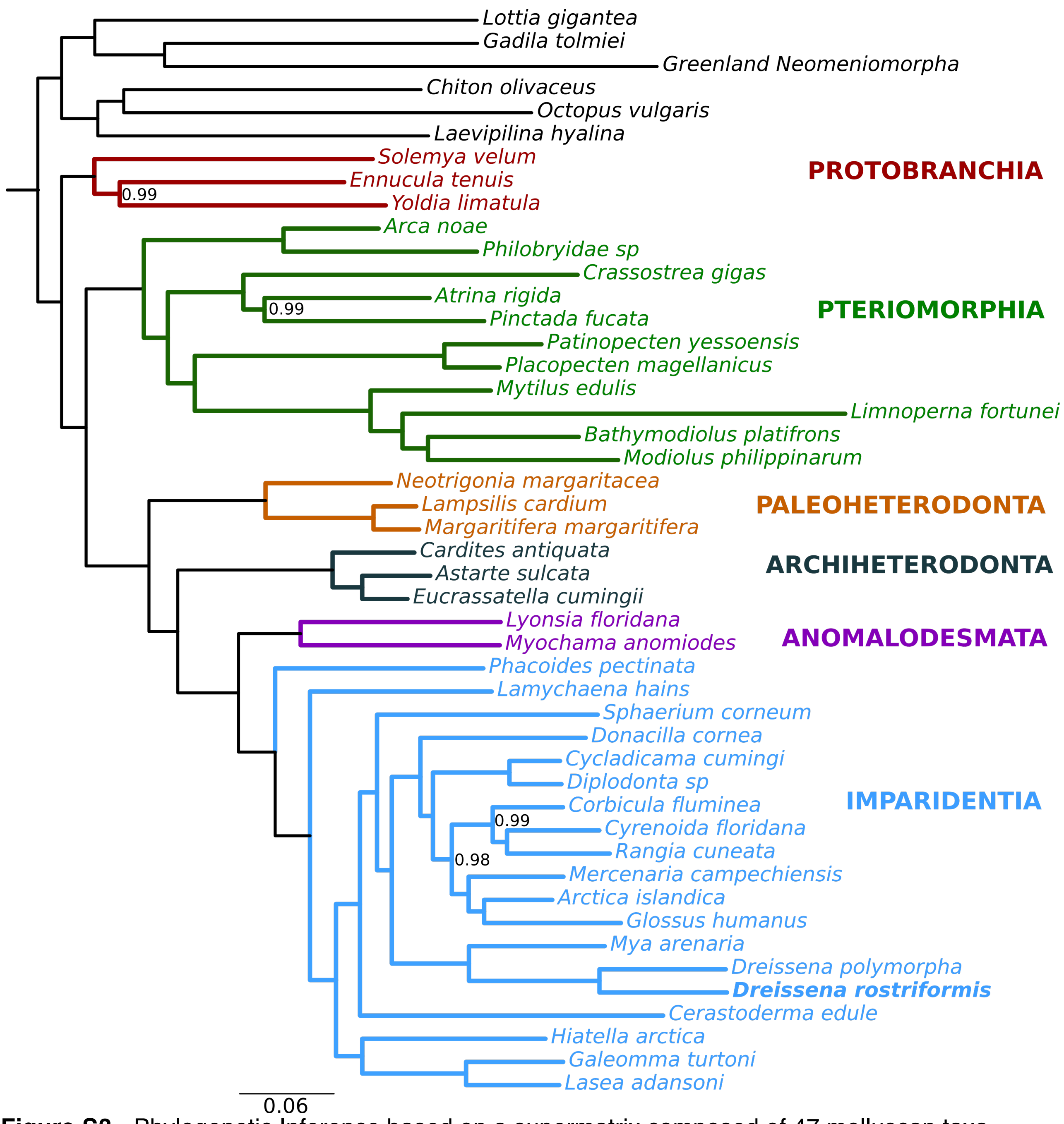

**Figure S3** - Phylogenetic Inference based on a supermatrix composed of 47 molluscan taxa (sequence length: 231,708; 1,377 genes) using FastTree version 2.1.10, under LG model of amino acid substitution and 100 bootstrap replicates. The 1,377 predefined orthogroups, originally composed of 40 molluscan taxa (*6*), were extended with seven new taxa using HaMStr software. The different colours in the tree correspond to different bivalve lineages, and bootstrap support values lower than 100 are indicated. Freshwater species are marked with a *.

**SM 3 - DNA Library preparation and sequencing**

**SM 3.1 - Genomic DNA extraction**

A single male *Dreissena rostriformis* selected for DNA extraction was collected from the Danube river in Vienna, Austria (48°14'45.9"N 16°23'38.0"E). Sample preparation and extraction proceeded as follows:

A. Sample preparation

A mature male mussel was kept in an aquarium in the laboratory and starved for two weeks in order to flush the digestive tract and to minimise contamination of the resulting DNA. The shell of the animal was then thoroughly cleaned using needle-nosed forceps under a dissecting microscope to remove mussel-associated annelids, sponges and macroscopic algae. The shell was briefly sprayed with 70% ethanol and dried before being placed in a clean Schott bottle with 100 mL of tap water and a 1X solution of antibiotic-antimycotic (Gibco 15240062). After 24 and 48 hours respectively, this solution was replaced with fresh antibiotic-antimycotic in tap water. 72 hours after the antibiotic-antimycotic treatment, the animal was cleaned and dissected in preparation for DNA extraction.

B. Dissection, DNA extraction and purification

The whole *Dreissena* was removed from its shell using a scalpel, sliced in to three parts and incubated in 10X weight/volume lysis buffer (*12*) on a gently rotating shaker table for approximately 25 hours at 50ºC. DNA was extracted by adding an equal volume of Phenol:Chloroform:Isoamyl alcohol (PCI, Sigma Aldrich P2069) to the lysate and mixing by gently rotating the tube for an hour until an emulsion formed. Phase separation involved centrifugation of the lysate:PCI solution at 5000g at room temperature followed by transfer of the resultant aqueous phase to a fresh tube using a wide-bore pipette. PCI treatment and phase separation were repeated five times.

To precipitate the DNA, 0.2X volume of 10M ammonium acetate and 2X volume of 96% ethanol were added to the extracted aqueous phase and incubated at room temperature overnight. The precipitated DNA was removed from the solution using a sterile glass hook and washed with 70% ethanol. Washing was repeated three times followed by air-drying of the resultant DNA pellet. Elution of the DNA pellet in TE buffer took approximately 36 hours at 55°C.

**SM 3.2 - Sequencing strategy**

Library preparation was outsourced to Eurofins Genomics, Ebersberg, Germany. In total, four shotgun and three mate pair libraries were produced. These libraries were pooled and sequenced over four lanes on an Illumina HiSeq2500 using v4 chemistry in high-output mode with 2 x 125 bp paired-end reads (Table S1).

**Table S1 -** Genomic DNA library data

| Shotgun library name | Lane | Insert size (bp) | Sequenced base pairs (Mbp) | Genome coverage |
|---|---|---|---|---|
| SG300_ACAGTG | 7 | 300 | 10,626 | 6.6x |
| SG300_GTGAAA | 7 | 300 | 13,498 | 8.4x |
| SG300_ACAGTG | 8 | 300 | 10,846 | 6.8x |
| SG300_GTGAAA | 8 | 300 | 13,654 | 8.5x |
| SG300_ACAGTG | 1 | 300 | 25,484 | 15.9x |
| SG300_GTGAAA | 1 | 300 | 32,257 | 20.2x |
| PCRfree550_CTTGTA | 3 | 550 | 20,617 | 12.9x |
| PCRfree550_GCCAAT | 3 | 550 | 20,795 | 13.0x |
| TOTAL | | | 147,777 | 92.4x |

| Mate pair library name | | Insert size (Kbp) | Read pairs | Genome fragment coverage |
|---|---|---|---|---|
| DreissenaDVA_LJD_3kb | 1,2 | 2.06 | 12,242,722 | 15.8x |
| DreissenaDVA_LJD_8kb | 1,2 | 6.5 | 24,885,918 | 101.6x |
| DreissenaDVA_LJD_20kb | 1,2 | 19.1 | 17,662,393 | 210.4x |

**SM 3.3 - Data pre-processing**

Proprietary read processing of the long jumping distance (LJD) libraries including quality and adaptor trimming was performed by Eurofins Genomics. Quality and adaptor trimming of the shotgun libraries was performed with trimmomatic (v0.35) (*13*) and library quality was assessed with FastQC (www.bioinformatics.babraham.ac.uk/projects/fastqc/).

**SM 3.4 - Estimation of genome size and heterozygosity**

Kmer assessment was performed with jellyfish (*14*) on all the libraries used in the assembly:

```
jellyfish count -t 16 -C -m 21 -s 8G *.bed -o reads.jf
jellyfish histo t 10 reads.jf > reads.histo
```

This was uploaded to GenomeScope (*15*) to estimate genome size the level of heterozygosity (Table S2)

**Table S2 -** GenomeScope assessment of genome assembly

| Property | min | max |
|---|---|---|
| Heterozygosity | 2.38% | 2.39% |
| Genome Haploid Length | 1,336,457,158 bp | 1,336,856,190 bp |
| Genome Repeat Length | 614,604,288 bp | 614,787,793 bp |
| Genome Unique Length | 721,852,870 bp | 722,068,397 bp |
| Model Fit | 94.06% | 97.80% |
| Read Error Rate | 0.02% | 0.02% |

The genome size estimated by GenomeScope differed from other methods. A previous report that used Feulgen image analysis densitometry (*16*) found a genome size for the closely related *D. polymorpha* of 1.7 pg which, when converted using the formula:

number of base pairs = mass in pg x 0.978 x 10^9

equates to a genome size of 1.66 Gb.

Using GCE (v1.0.0) with a 19mer kmer graph (*17*):

```
gce -f reads.histo -g 117478492290 -c 75 -H 1 -m 1 -D 8
```

results in an estimated genome size of 1.56 Gb.

Manual calculation using the formula:

N = (M*L)/(L-K+1), Genome_size = T/N,
where N: Depth, M: Homo Kmer peak, K: Kmer-size, L: avg readlength,
T: Total bases

gives a genome size estimate of 1.58 Gb.

At k32, a high heterozygous peak at 31x and a lower homozygous peak at 63x is indicative of the high level of heterozygosity estimated by GenomeScope (Fig. S4).

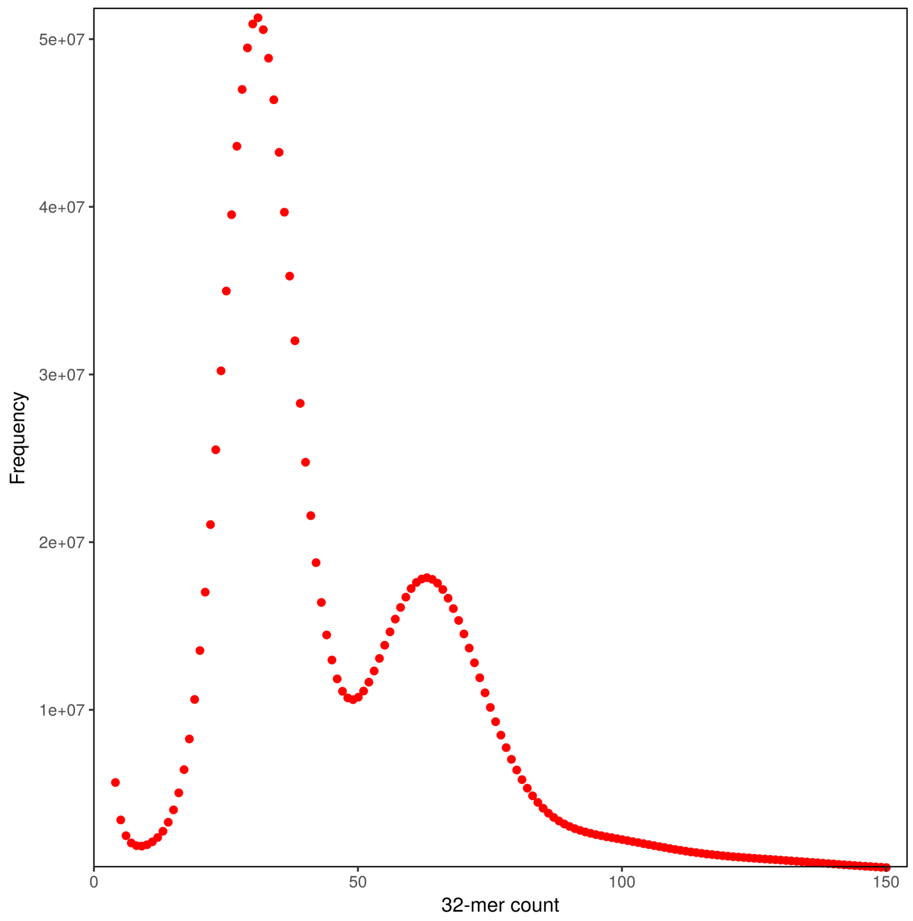

**Figure S4 -** 32-mer depth distribution.

### SM 4 - Genome assembly and quality control

#### SM 4.1 - Genome assembly

Genome contig assembly, scaffolding and gap-closing were performed with Platanus v1.2.4 (*18*). Assembly, scaffolding and gap-closing were performed on the SGI Altix Ultra Violet 1000 located at the Johannes Kepler University, Linz, Austria. The following options were used for contig assembly, scaffolding and gap-closing respectively:

-u 0.3 -d 0.3 -k 32 -t 96

-t 96 -u 0.3

-t 96

Following Platanus assembly, heterozygosity was reduced using the Redundans pipeline (v0.13a) (*19*). This was performed on the Life Science Compute Cluster (CUBE) located at the University of Vienna, Austria. Redundans options selected were:

identity=0.51

minLength=200

overlap=0.66

**SM 4.2 - Genome quality assessment**

Quast was used to assess properties of the genome assembly (Table S3).

**Table S3 -** Quast genome assessment

| Assembly | Dpo_1.0 |
|---|---|
| # contigs (>= 0 bp) | 18,504 |
| # contigs (>= 1000 bp) | 18,504 |
| # contigs (>= 5000 bp) | 18,504 |
| # contigs (>= 10000 bp) | 17,432 |
| # contigs (>= 25000 bp) | 10,989 |
| # contigs (>= 50000 bp) | 7,110 |
| Total length (>= 0 bp) | 1,241,502,953 |
| Total length (>= 1000 bp) | 1,241,502,953 |
| Total length (>= 5000 bp) | 1,241,502,953 |
| Total length (>= 10000 bp) | 1,231,413,561 |
| Total length (>= 25000 bp) | 1,128,185,434 |
| Total length (>= 50000 bp) | 989,540,184 |
| # contigs | 18,504 |
| Largest contig | 1,148,001 |
| Total length | 1,241,502,953 |
| GC (%) | 34.88 |
| N50 | 131,410 |
| N75 | 61,075 |
| L50 | 2,627 |
| L75 | 6,055 |
| # N's per 100 kbp | 5,074.71 |

The difference between the assembly length (1.24 Gbp) and the predicted length (~1.6 Gbp) is likely due to the inability of Platanus to assemble long highly repetitive regions. As such, the missing sequences are likely to be highly repetitive and gene-poor. Similar results were reported for the scallop genome, *Patinopecten yessoensis* (*20*).

**SM 4.3 - Read re-mapping**

The shotgun libraries were mapped back to the completed genome assembly with Bowtie 2 (*21*) to assess assembly integrity. In total, 94.45% of the shotgun reads were successfully mapped back to the genome (Table S4).

**Table S4 -** Library re-mapping to completed genome assembly

| | SG300_ACAGTG_L7 | SG300_GTGAAA_L7 | SG300_ACAGTG_L8 | SG300_GTGAAA_L8 | SG300_ACAGTG_L1 | SG300_GTGAAA_L1 | PCRfree550_CTTGTA_L3 | PCRfree550_GCCAAT_L3 |
|---|---|---|---|---|---|---|---|---|
| Total paired reads | 40183468 | 51913888 | 41213936 | 52750257 | 97693522 | 124792269 | 78371937 | 79934150 |
| Read pairs aligned concordantly 1 time | 83.80% | 83.90% | 84.60% | 84.70% | 85.20% | 85.50% | 54.60% | 54.00% |
| Read pairs aligned dicordantly 1 time | 8.20% | 8.20% | 7.80% | 7.80% | 7.70% | 7.60% | 32.10% | 32.70% |
| Mates aligned 1 time | 1.60% | 1.50% | 1.60% | 1.50% | 1.50% | 1.40% | 2.90% | 2.90% |
| Mates aligned >1 time | 55.00% | 0.50% | 0.50% | 0.50% | 0.50% | 0.50% | 4.40% | 4.50% |
| Overall alignment rate | 94.10% | 94.10% | 94.50% | 94.50% | 94.90% | 95.00% | 94.00% | 94.00% |

### SM 4.4 - Genomic contamination

To assess whether any bacterial scaffolds were included in the assembly, all the genes located on scaffolds with G+C content two standard deviations higher or lower than the mean G+C content (Fig. S5) were inspected using BLASTP (v.2.6.0) against the UNIREF90 database (release 2018_03). Of the 903 scaffolds inspected, 434 encoded at least one gene model and only 84 encoded at least two gene models. Inspection of the 194 gene models located on these 84 scaffolds against the UNIRE90 database identified a single gene with a top hit to a bacterial sequence (*Gene.957*, UniRef90_*A0A0T6LNL7*). Further inspection of this sequence against the NCBI nr database identified sequence similarity with various molluscs, cnidarians and vertebrates in addition to bacterial sequences leading to an ambiguous homology determination. The scaffold hosting *Gene.957* (scaffold10118) also encodes two other genes (*Gene.956*, *Gene.960*), which together with *Gene.957* are all multi-exonic. BLASTP of *Gene.956* against the NCBI nr database identifies sequence similarity with replication factor C subunit 5-like sequences from various molluscs. No sequence similarity was identified for *Gene.960*.

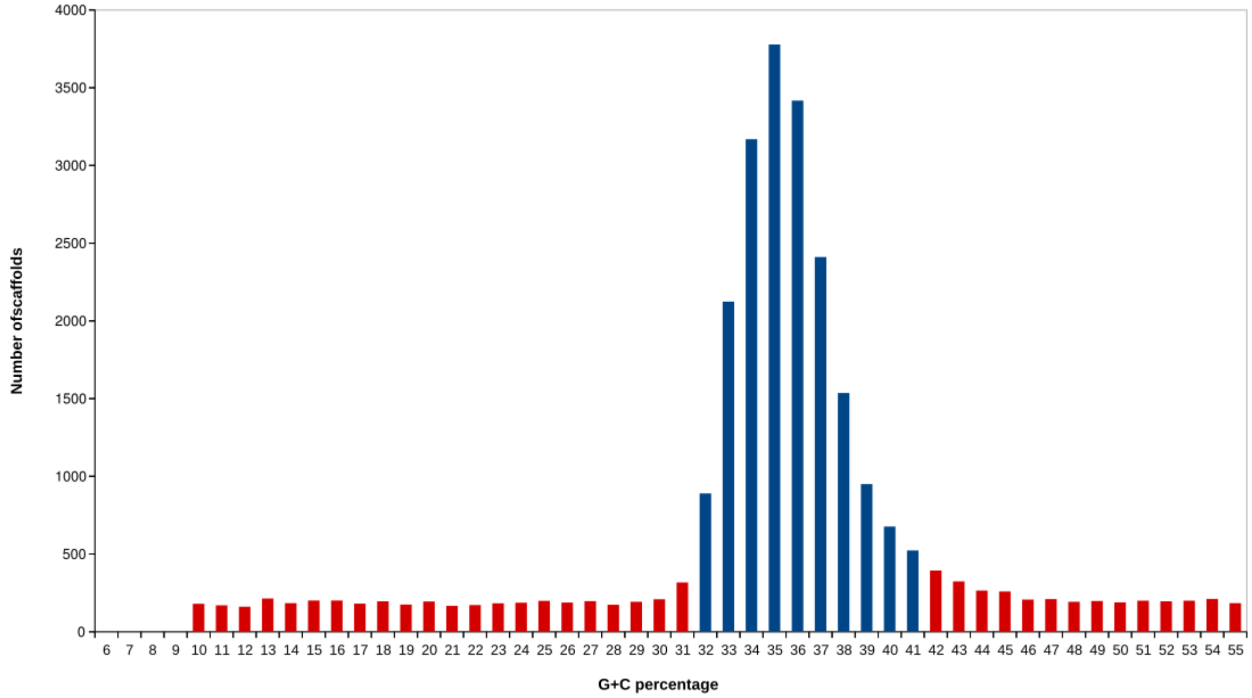

**Figure S5 -** G+C content of the 18,505 scaffolds that make up the quagga mussel assembly binned to 1%. Those with a G+C content higher or lower than two standard deviations of the mean G+C content (36.3%) are coloured in red while the remainder are coloured in blue.

To further assess genome contamination, a blobplot (*22*) was constructed by mapping the eight libraries mentioned in Table S4 back to the assembly and by using MEGABLAST to obtain taxids for the genomic scaffolds (Figs. S6, S7). No bacterial contamination was identified however some scaffolds were designated non-molluscan taxids, the most abundant of which were 'chordate' (1.3% of scaffolds).

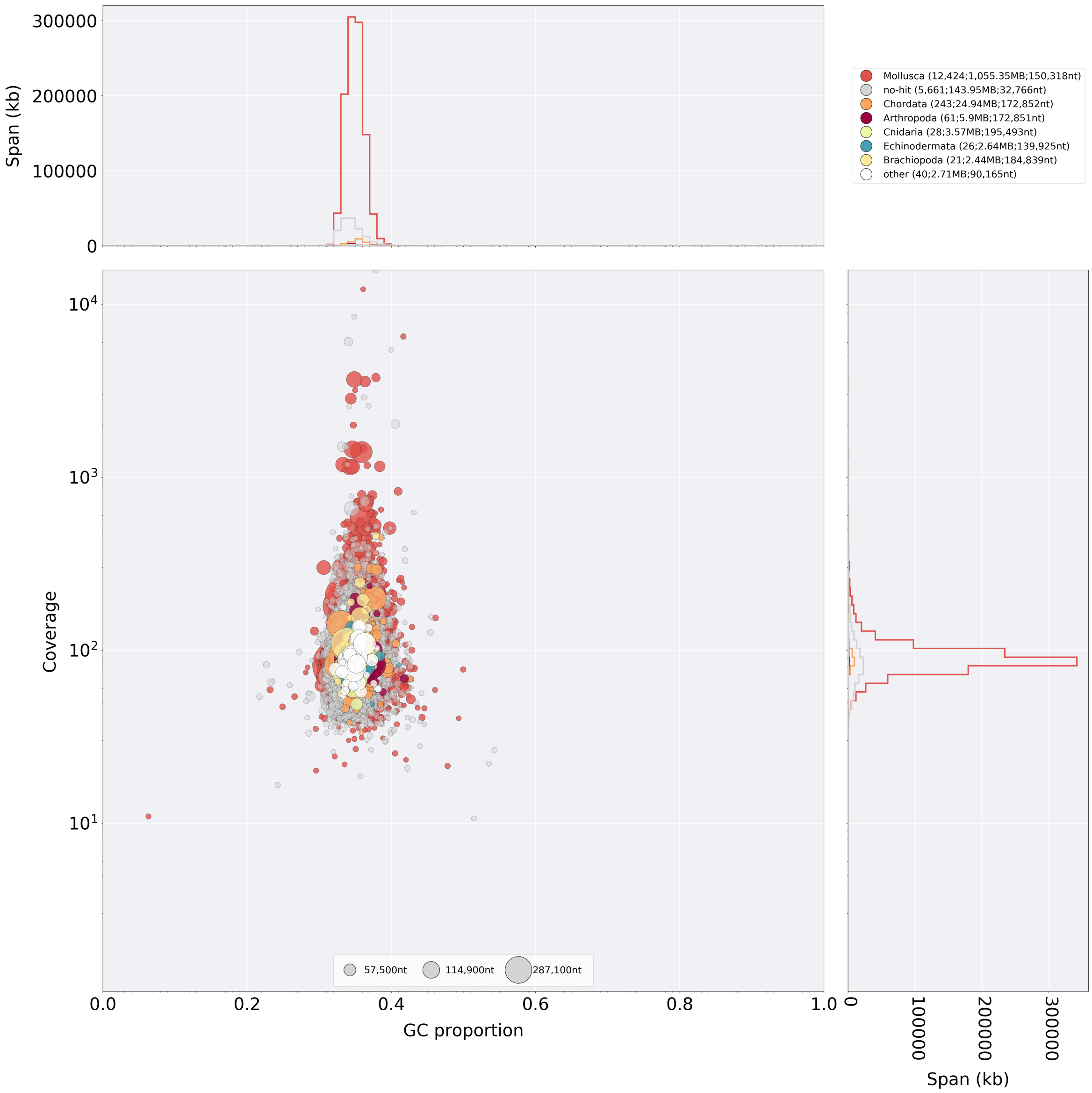

**Figure S6 -** Blobplot of genome assembly. Determination of potential contamination with BlobTools involves plotting assembled scaffolds by their coverage and GC proportion to identify groups of scaffolds with distinct properties. The absence of distinct blobs which are typical of contaminated assemblies suggests a lack of contamination.

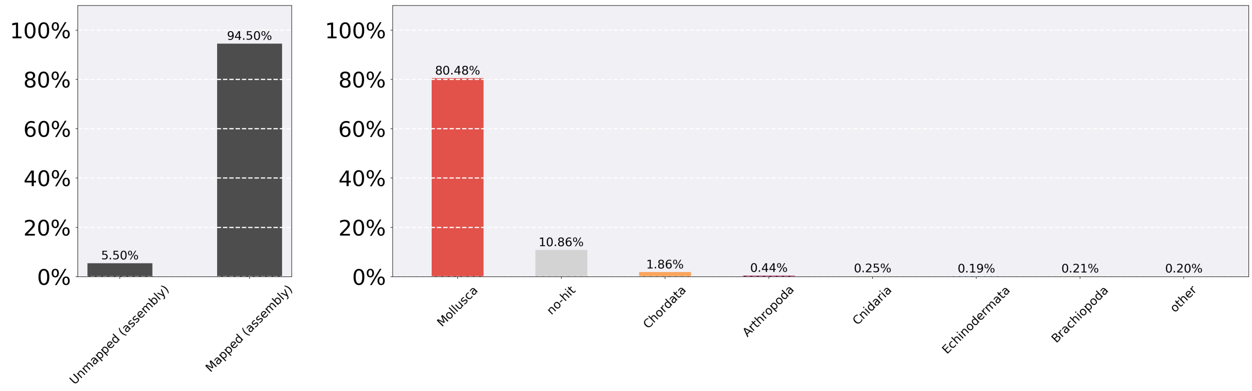

**Figure S7 -** BlobTools ReadCovPlot output. In total, 91% of reads mapped to scaffolds that were annotated as either molluscan or unknown. The majority of the remaining scaffolds were determined to be chordate-like based on the output of MEGABLAST. No significant proportion of scaffolds was determined to be microbial.

To confirm the source of these scaffolds, the proteins encoded by genes (see SM 5) hosted by the 243 chordate-like scaffolds were BLASTed against the NCBI nr database with an e-value cutoff of 1e-6 and allowing for 500 hits. The species with the best hit as determined by the bit score was then determined resulting in just 50 scaffolds hosting at least one chordate-like gene. Of these 50 scaffolds, only three did not host a non-chordate like gene in addition to a chordate-like gene. As the BLASTP results did not produce any consistent pattern to suggest contamination by a particular chordate taxon and because of the total length of scaffolds hosting only chordate-like genes equated to only 0.004% of the total assembly length (53,307 bp), it was decided not to discard any of these scaffolds from the assembly.

### SM 5 - Genome annotation

#### SM 5.1 - Repeat annotation

Construction of a RepeatModeler (*23*) library uncovered 1842 elements, 1428 of which were unknown. In total 31.88% of the genome was masked by RepeatMasker, the majority of which (24.2%) are unclassified (Table S5).

**Table S5 -** RepeatMasker output.

```
==================================================
file name: Dpo_1.0.fa
sequences:          18504
total length: 1241502953 bp  (1178602863 bp excl N/X-runs)
GC level:         34.88 %
bases masked:  395771212 bp ( 31.88 %)
==================================================
               number of      length   percentage
               elements*    occupied  of sequence
--------------------------------------------------
SINEs:            44757      8501253 bp    0.68 %
      ALUs            0            0 bp    0.00 %
      MIRs         6527      1293685 bp    0.10 %

LINEs:            79828     34271882 bp    2.76 %
      LINE1        1749       466299 bp    0.04 %
      LINE2        9068      3715883 bp    0.30 %
      L3/CR1       4548      1989511 bp    0.16 %

LTR elements:     17697      5283270 bp    0.43 %
      ERVL            0            0 bp    0.00 %
      ERVL-MaLRs      0            0 bp    0.00 %
      ERV_classI    345       131396 bp    0.01 %
      ERV_classII     0            0 bp    0.00 %

DNA elements:    138531     30530243 bp    2.46 %
     hAT-Charlie   1382       149247 bp    0.01 %
     TcMar-Tigger     0            0 bp    0.00 %

Unclassified:    1511489    300414069 bp    24.20 %

Total interspersed repeats:379000717 bp   30.53 %

Small RNA:         1789       474488 bp    0.04 %

Satellites:         730       267606 bp    0.02 %
Simple repeats:  215638     16335930 bp    1.32 %
Low complexity:   23582      1116226 bp    0.09 %
==================================================
```

### SM 5.2 - RNA preparation and sequencing

In order to annotate protein coding genes, four developmental RNA seq libraries were constructed. To produce developmental material and to maximise the number of expressed genes, quagga mussels and closely related zebra mussels (Fig. S1), were collected from the Danube river in Vienna, Austria, and were induced to spawn through immersion for five minutes in a solution of 0.5 mM serotonin (Sigma-Aldrich H9523) in filtered river water (FRW). Following serotonin treatment, the adults were separated into individual glass dishes until spawning occurred. Eggs were fertilised through the introduction of sperm for thirty minutes, after which the fertilised eggs were washed thoroughly with FRW to remove excess sperm. Pooled embryos (quagga and zebra) were allowed to develop at room temperature (~23°C) until they had reached the desired stage of development. Samples of gastrulas (approximately 5-6 hours post fertilisation), trochophores (approximately 11-12 hours post fertilisation) and early veligers (approximately 36 hours post fertilisation) were sampled and stored in RNAlater at 4°C overnight before being transferred to a freezer at -20°C. In addition, a single juvenile zebra mussel (*Dreissena polymorpha*) approximately 3 mm in shell length was dissected from its shell and stored in RNAlater for the juvenile sample. RNA extractions were conducted using the Qiagen RNA kit as per the instructions. RNA samples were DNAse treated.

RNA samples were sent to the Vienna Biocenter Core Facility (VBCF) for library construction and sequencing. For all samples, RNA-seq libraries were constructed with the Lexogen SENSE mRNA-Seq Library Prep Kit V2 and were sequenced on an Illumina Hi-Seq 2500 generating paired-end, stranded 125 bp libraries (Table S6).

**Table S6 -** Transcriptome assembly RNAseq data summary

| Sample | Description | Reads (paired end) | Size (Mb) |
|---|---|---|---|
| Gastrula | pooled gastrulas | 96,575,472 | 12,071 |
| Trochophore | pooled trochophores | 105,021,022 | 13,128 |
| Early Veliger | pooled veligers | 113,419,094 | 14,177 |
| Juvenile | single 3mm mussel | 95,097,238 | 11,887 |

#### SM 5.3 - *De novo* transcriptome assembly

Following Lexogen's instructions, the first library of the pair was trimmed to remove the first nine nucleotides of each read and the second library of the pair was trimmed to remove the first six nucleotides of each read. Adapters and low quality sequence were removed with trimmomatic v 0.35 (*13*). Five transcriptomes were built for each library with Binpacker (*24*) using k23, k25, k27, k29 and k32. Individual kmer assemblies were then merged with Velvet (*25*) and de-duplicated with Dedupe (*26*). Open reading frames (ORFs) were predicted with Transdecoder v 3.0.0 using the –single_best_only option (*27*). All four transcriptomes were then concatenated and de-duplicated with Dedupe and UCLUST v1.2.22q (*28*). Transdecoder was used once more to predict ORFs from this concatenated assembly. 65% of the 169875 transcripts (109955) were mapped back to the assembled genome with GMAP (v 2016.01.21) (*29*) allowing for 75% minimum trimmed coverage and 50% minimum identity. The low percentage of mapped transcripts was likely due to the high level of heterozygosity expected in quagga mussel populations and due to the existence of some zebra mussel specific transcripts.

**SM 5.4 - Reference-based transcriptome assembly**

A reference-based transcriptome was produced with the same four RNA-seq libraries used in the *de novo* assembly. Each of the trimmed libraries were mapped against the reference genome using STAR aligner v2.5.0a (*30*) and then assembled with StringTie v1.3.3b (*31*). Assemblies were merged with the StringTie merge function and ORFs predicted with TransDecoder. The resulting transcriptome consisted of 60,557 coding transcripts.

**SM 5.5 - *Ab initio g*ene prediction**

A training set of 380 genes was created for Augustus (*32*) from the *de novo* transcriptome assembly. The training gene set all mapped to the genome with 100% accuracy, contained at least 3 exons, had start and stop codons and had homology to a sequence in either the Pfam, uniref90 or CDD databases. Using the training dataset, Augustus predicted 72,428 transcripts. Gene prediction was also conducted with SNAP (*33*) producing a set of 113,706 coding transcripts.

**SM 5.6 - Homology-based gene prediction**

The complete annotated protein complement of five species (*Crassostrea gigas*, *Octopus bimaculoides*, *Lottia gigantea*, *Lingula anatina* and *Drosophila melanogaster*) were aligned to the *Dreissena* genome with TBLASTN (E-value ≤ 1e-5). These were then passed to GeneWise v2.4.1 (*34*) to produce accurate spliced alignments. In total 102,299 spliced alignments were identified.

**SM 5.7 - Gene model evaluation**

The output of the *de novo* transcriptome assembly, the reference based transcriptome assembly, the two gene prediction methods (Augustus and SNAP) and the homology-based gene prediction were used as input for EvidenceModeler (EVM; *35*). The weights for EVM were as per Table S7.

**Table S7 -** EvidenceModeler inputs and weights

| Evidence Type | Details | Weight |
|---|---|---|
| OTHER_PREDICTION | Gmap mapped Binpacker de novo transcripts | 20 |
| OTHER_PREDICTION | Stringtie reference based transcripts | 20 |
| PROTEIN | GeneWise homology based transcripts | 10 |
| ABINITIO_PREDICTION | Augustus gene predictions | 1 |
| ABINITIO_PREDICTION | SNAP gene predictions | 1 |

The 99,522 gene models produced by EVM were then filtered to only include those that have homology to a sequence in either the Pfam, uniref90 or CDD databases or for which there is evidence of expression in one of the developmental RNA-seq databases as assessed by Kallisto (SM 6.2; *36*). Gene models that overlapped with repetitive sequences as assessed by RepeatMasker (SM 5.1; *23*) for at least 50% of their length were also excluded. The final transcriptome consisted of 37,681 coding genes which included 95% of the metazoan BUSCO v2.0 genes when run in protein mode (Table S8; Fig. S8; *37*). The genome sequencing, assembly and annotation pipeline are summarised in Fig. S9.

**Table S8 -** Quagga mussel transcriptome data

| **Transcriptome data** | |
|---|---|
| Number of gene models | 38,084 |
| Number of reconstructed bases | 43,727,907 |
| GC content | 48.10% |
| BUSCOs complete | 83.30% |
| BUSCOs complete, single copy | 80.20% |
| BUSCOs complete duplicated | 3.10% |
| BUSCOs fragmented | 11.70% |
| BUSCOs missing | 5.00% |
| BUSCOs identified (complete plus fragmented) | 95.00% |
| Average number of exons | 5.9 |
| Average exon length | 195 |
| Average intron length | 1,340 |
| Homology support (total) | 32,708 |
| Homology support (Pfam) | 25,850 |
| Homology support (uniref90) | 32,283 |
| Homology support (CDD) | 25,666 |

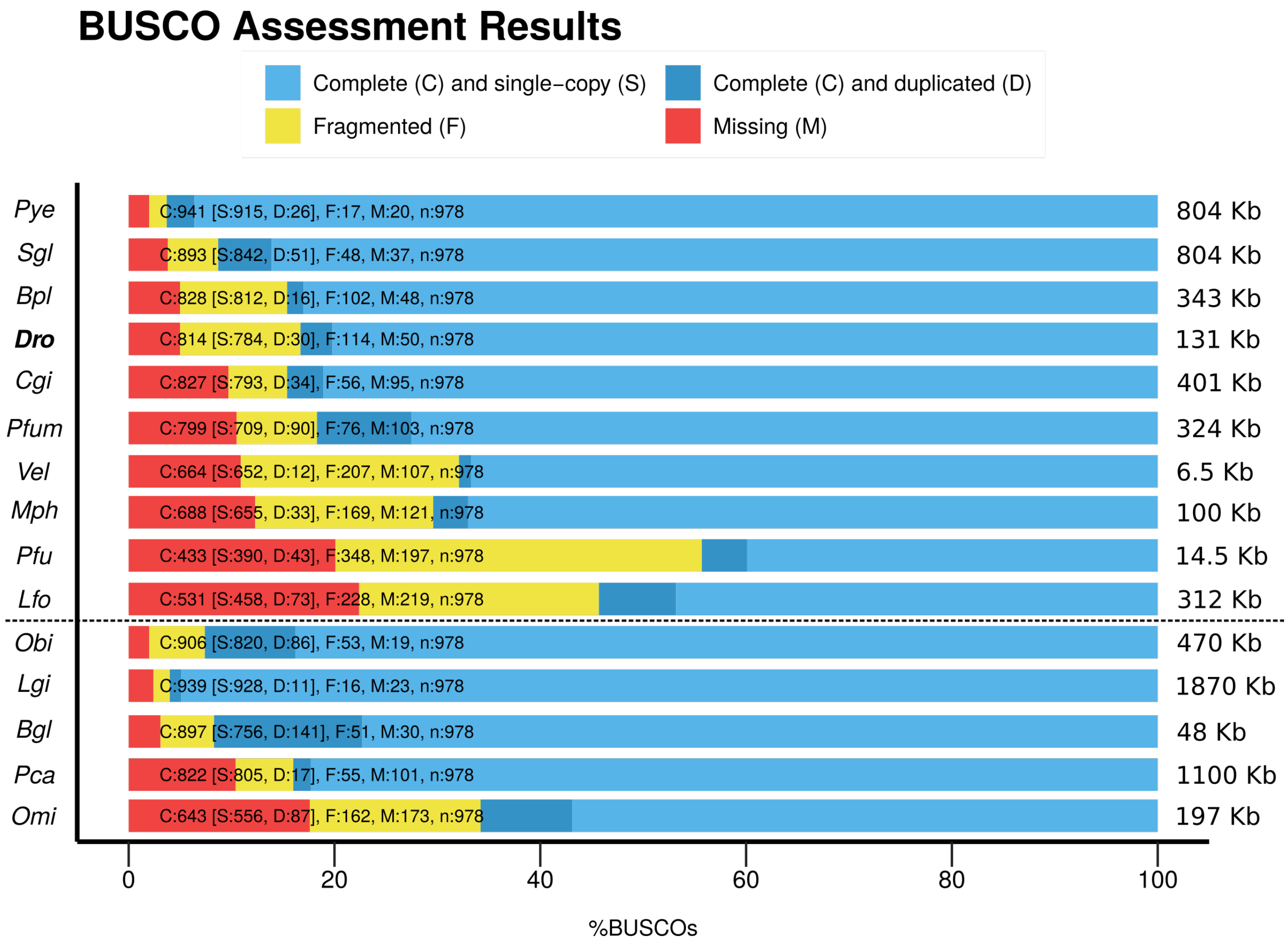

**Figure S8** - Comparative BUSCO scores for molluscan genomes. Above the dotted line are the BUSCO scores for bivalve species and below the line are the scores for other molluscan taxa. Species are ordered from lowest number of missing BUSCOs to highest with the scaffold N50 for each species displayed on the right. *Pye - Patinopecten yessoensis, Sgl - Saccostrea glomerata, Bpl - Bathymodiolus platifrons, Dro - Dreissena rostriformis, Cgi - Crassostrea gigas, Pfum - Pinctada fucata martensii, Vel - Venustaconcha ellipsiformis, Mph - Modiolus philippinarum, Pfu - Pinctada fucata, Lfo - Limnoperna fortunei, Obi - Octopus bimaculoides, Lgi - Lotti gigantea, Bgl - Biomphalaria glabrata, Pca - Pomacea canaliculata, Omi - Octopus minor.*

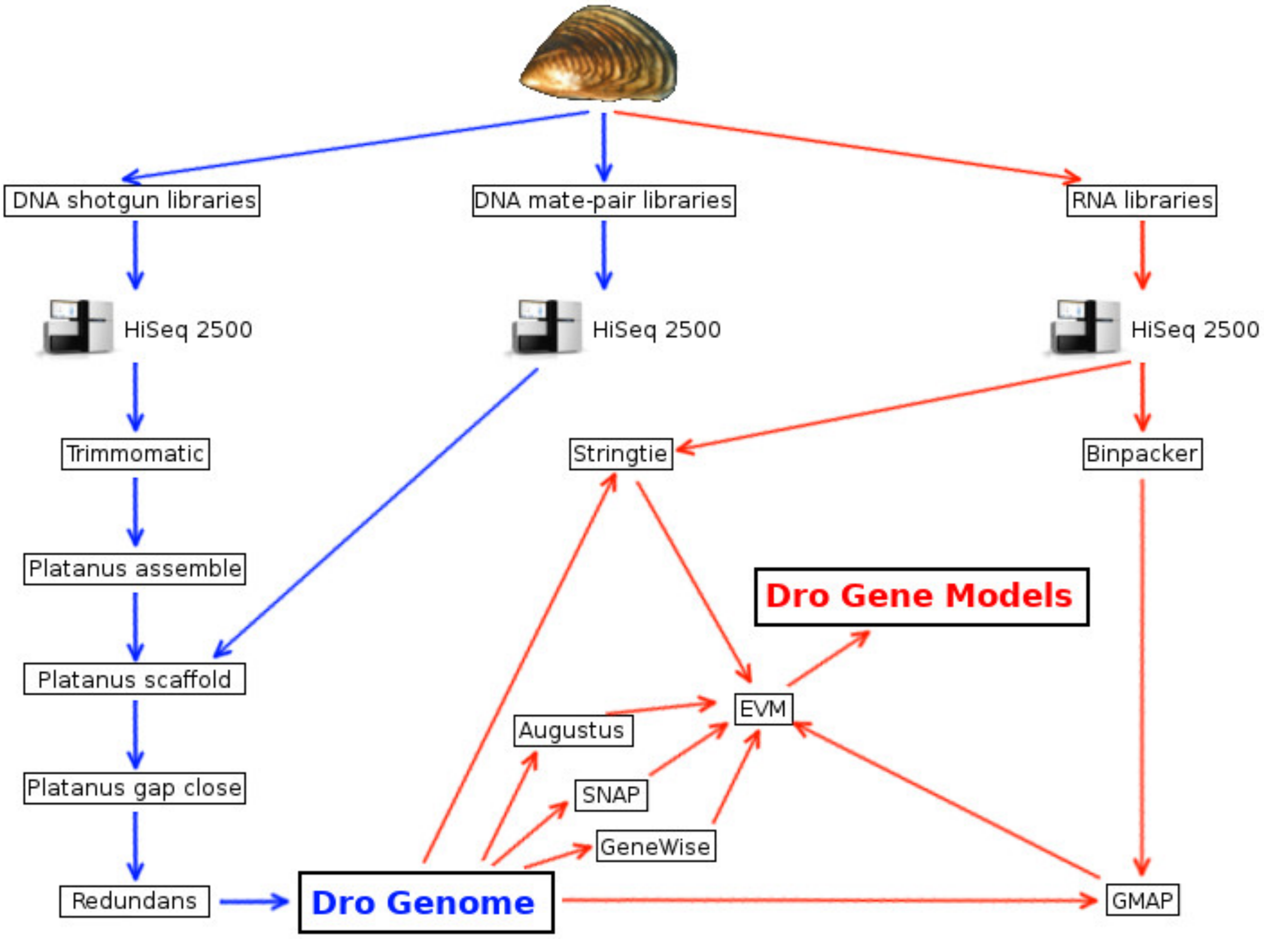

**Figure S9 -** Quagga mussel genome assembly and annotation pipeline.

### SM 6 - Identification and annotation of candidate osmoregulatory genes

#### SM 6.1 - Gene identification

To identify genes encoding proteins with known roles in osmoregulation, ionic homeostasis and excretion, the full set of *Dreissena* gene models were used to search against the KEGG database (*38*) using the KAAS search tool (*39*). We focused on genes encoding transmembrane proteins involved in one of five KEGG pathways: 1) vasopressin-regulated water reabsorption, 2) proximal-tubule bicarbonate reclamation, 3) collecting duct acid secretion, 4) aldosterone regulated sodium reabsorption and 5) endocrine and other factor calcium reabsorption (Figs. S10-S14).

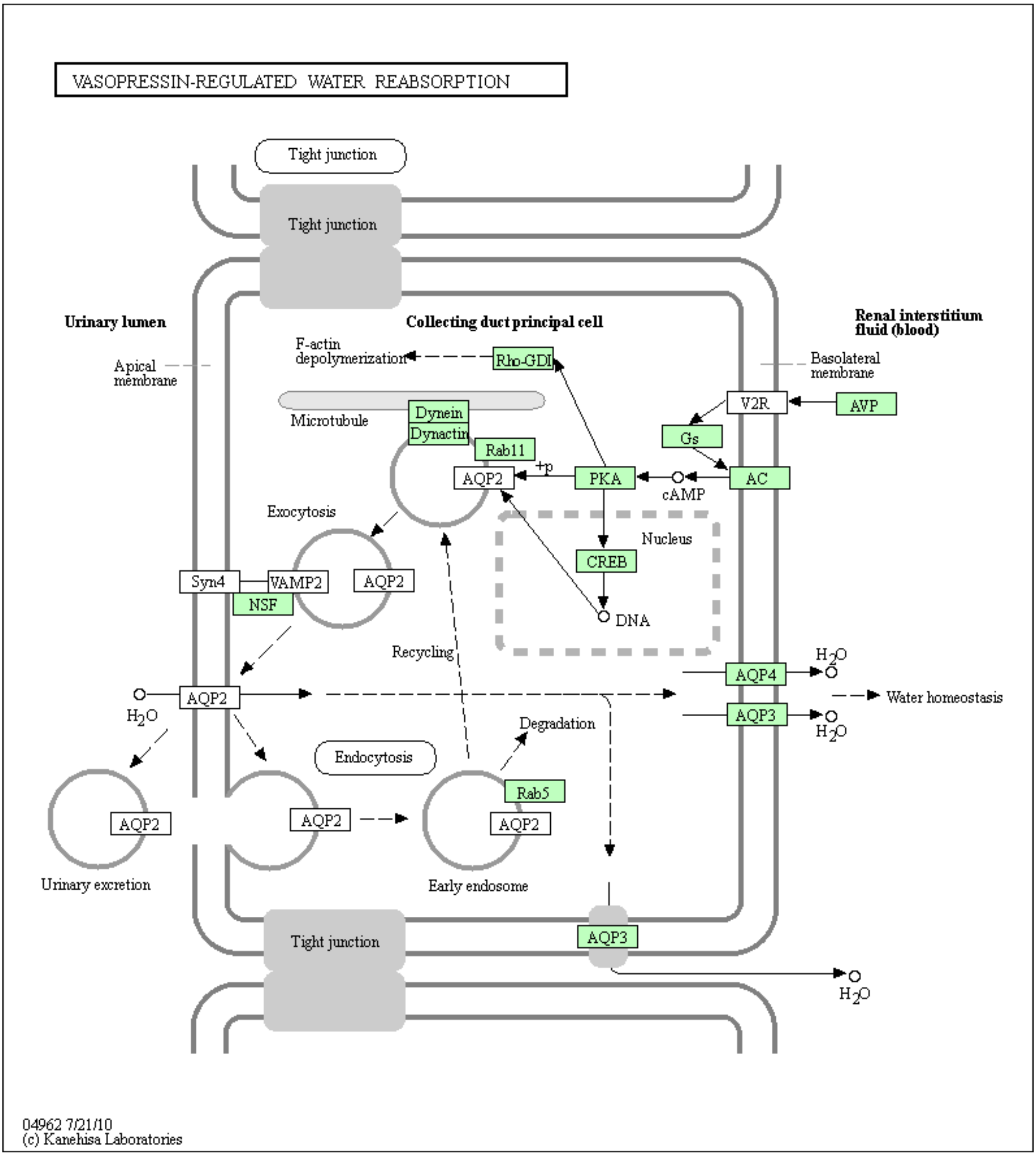

**Figure S10 -** Vasopressin-regulated water reabsorption KEGG pathway (*38*). Genes in green boxes were identified in the quagga mussel transcriptome using the KAAS search tool (*39*).

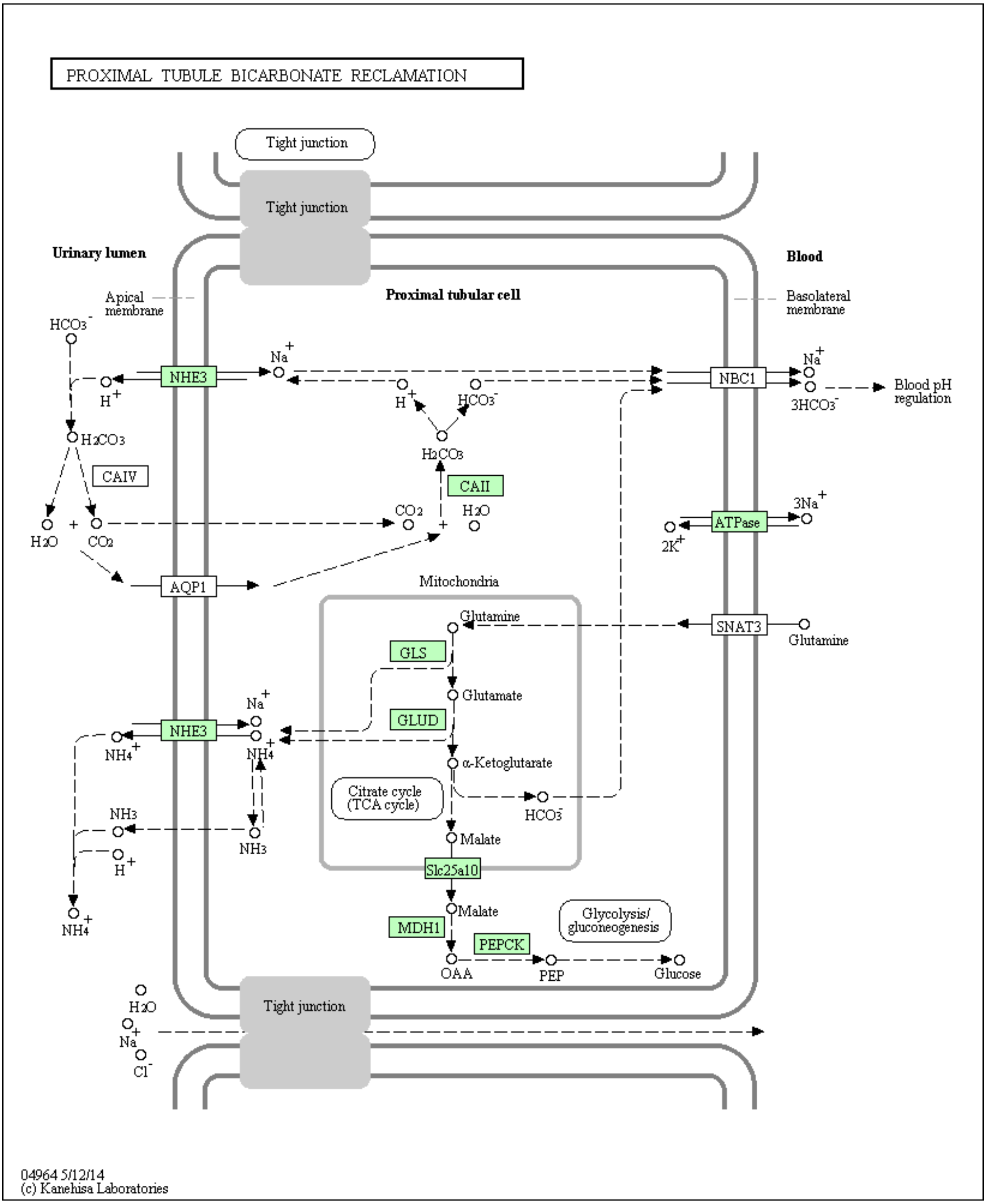

**Figure S11 -** Proximal-tubule bicarbonate reclamation KEGG pathway (*38*). Genes in green boxes were identified in the quagga mussel transcriptome using the KAAS search tool (*39*).

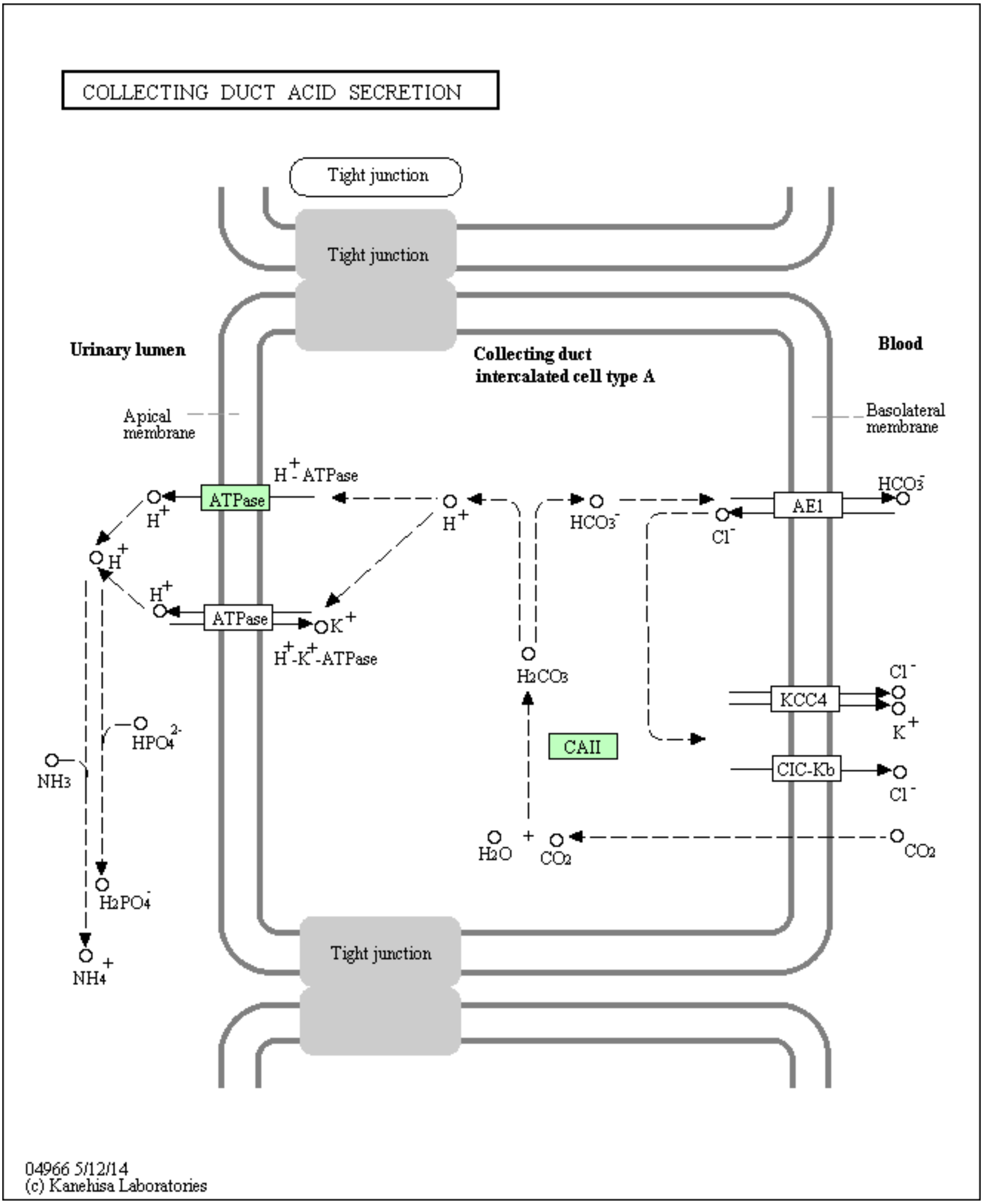

**Figure S12 -** Collecting duct acid secretion KEGG pathway (*38*). Genes in green boxes were identified in the quagga mussel transcriptome using the KAAS search tool (*39*).

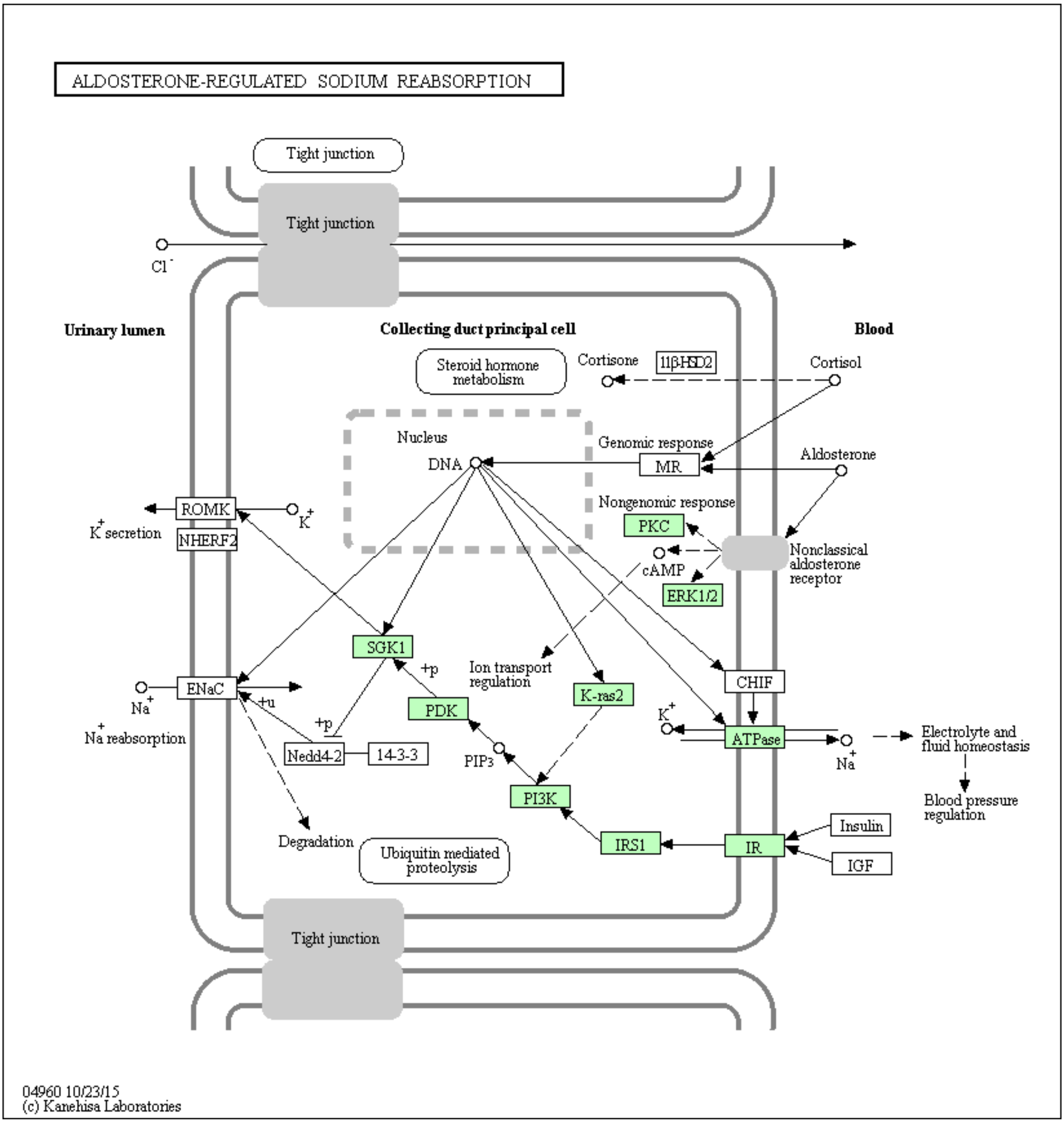

**Figure S13 -** Aldosterone regulated sodium reabsorption KEGG pathway (*38*). Genes in green boxes were identified in the quagga mussel transcriptome using the KAAS search tool (*39*).

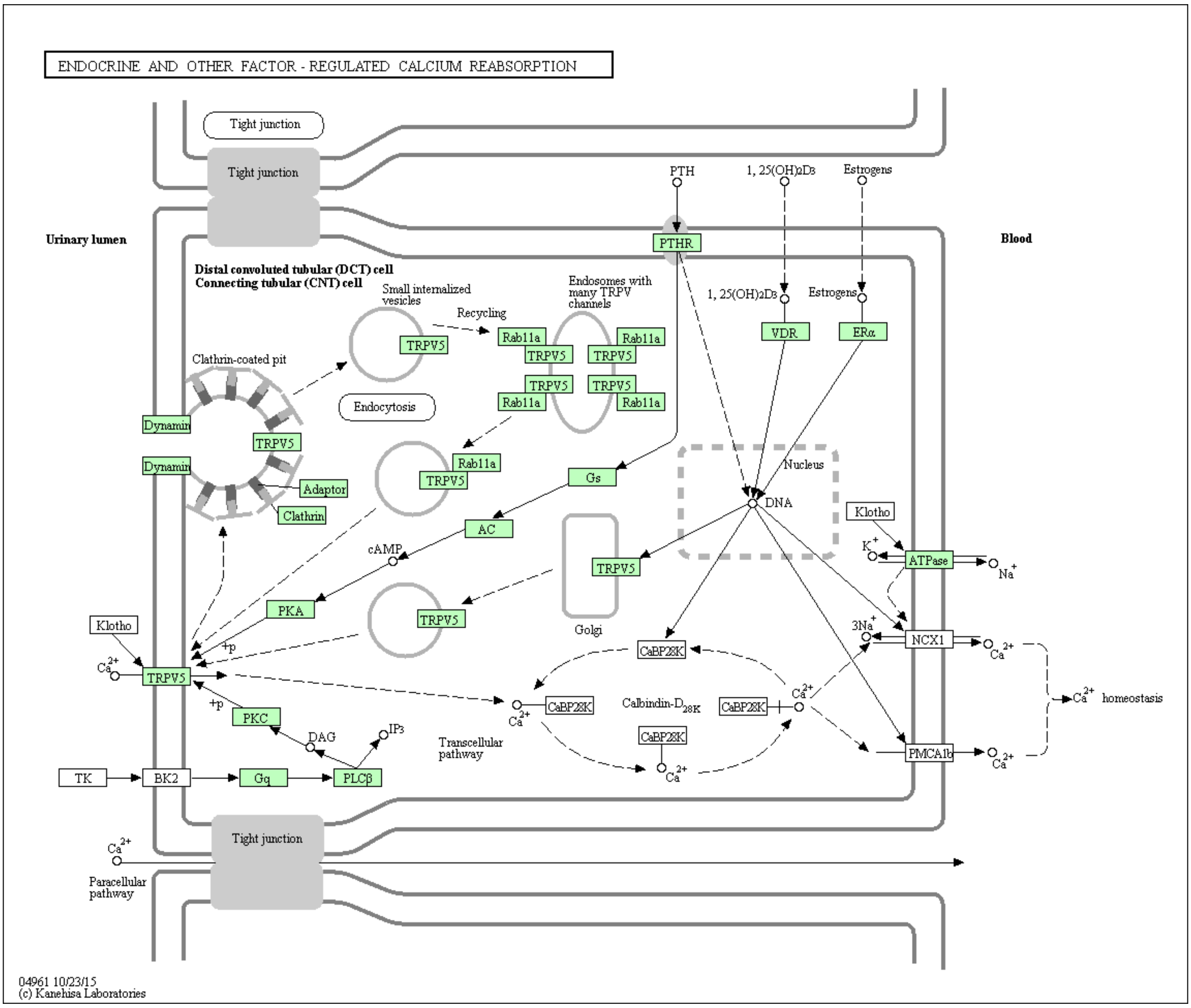

**Figure S14 -** Endocrine and other factor calcium reabsorption KEGG pathway (*38*). Genes in green boxes were identified in the quagga mussel transcriptome using the KAAS search tool (*39*).

Using HMMSEARCH from the HMMER3 package (*9*) with an e value of 1e-6, we identified 27 aquaporins (Pfam: PF00230.19), eight sodium/potassium ATPases (Pfam: PF00287.17), 13 sodium/hydrogen exchangers (NHE, Pfam: PF00999.20), eight hydrogen/carbonate co-transporters (Pfam: PF00955.20), 12 voltage gated chloride channels (PF00654.19), nine cation ATPases (Pfam: PF00689.20, PF13246.5, PF00690.25) and 17 hydrogen ATPases (Pfam: PF00006.24, PF02874.22, PF01813.16, PF03223.14, PF03179.14, PF01496.18, PF01992.15, PF01991.17) from the *Dreissena* transcriptome.

To determine which of these were highly expressed during early development, we produced 18 RNA-seq libraries from different developmental stages using the same protocol described for the four original RNA-seq libraries described in SM 5.2. These pooled samples were all barcoded and sequenced on a single lane of an Illumina HiSeq2500 (Table S9).

**Table S9 -** RNA-seq library data.

| Sample | Description | Reads (paired end) | Size (Mb) |
|---|---|---|---|
| 0hpf | Unfertilised eggs | 26,976,176 | 3,372 |
| 2hpf | 2-4 cell embryos | 15,272,738 | 1,909 |
| 4hpf | Gastrulas | 29,902,254 | 3,738 |
| 6hpf | Swimming gastrulas 1 | 28,373,154 | 3,547 |
| 8hpf | Swimming gastrulas 2 | 21,107,958 | 2,638 |
| 13hpf | Trochophores 1 | 23,421,090 | 2,928 |
| 18hpf | Trochophores 2 | 22,581,004 | 2,823 |
| 22hpf | Trochophores 3 | 26,543,548 | 3,318 |
| 23hpf | Trochophores 4 | 27,344,048 | 3,418 |
| 26hpf | Early veligers 1 | 30,418,842 | 3,802 |
| 27hpf | Early veligers 2 | 43,952,712 | 5,494 |
| 30hpf | Early veligers 3 | 33,454,982 | 4,182 |
| 36hpf | D-shaped veligers 1 | 25,858,654 | 3,232 |
| 48hpf | D-shaped veligers 2 | 37,566,002 | 4,696 |
| 54hpf | D-shaped veligers 3 | 41,105,390 | 5,138 |
| 60hpf | D-shaped veligers 4 | 39,914,750 | 4,989 |
| 72hpf | D-shaped veligers 5 | 39,113,060 | 4,889 |
| 84hpf | Late D-shaped veligers | 28,465,618 | 3,588 |

**SM 6.2 - Developmental expression dynamics**

All 18 libraries were pre-processed as per SM 5.3 before being pseudoaligned to the *Dreissena* transcriptome with Kallisto to determine the Transcripts Per Million (TPM) value for each gene (*36*). A comparison of the TPM values of the candidate osmoregulation, ionic homeostasis and excretion genes over the course of development in *Dreissena* and *Crassostrea* shows 1) no significant difference in the overall expression levels of this category of genes between the two species and 2) no enrichment for this category of genes in any particular developmental stage (Fig. S15). This indicates that the processes of osmoregulation, ionic homeostasis and excretion are important at each point of development, regardless of the environmental osmotic conditions and so a more nuanced approach is required to determine the specific molecular machinery required for embryonic osmoregulatory.

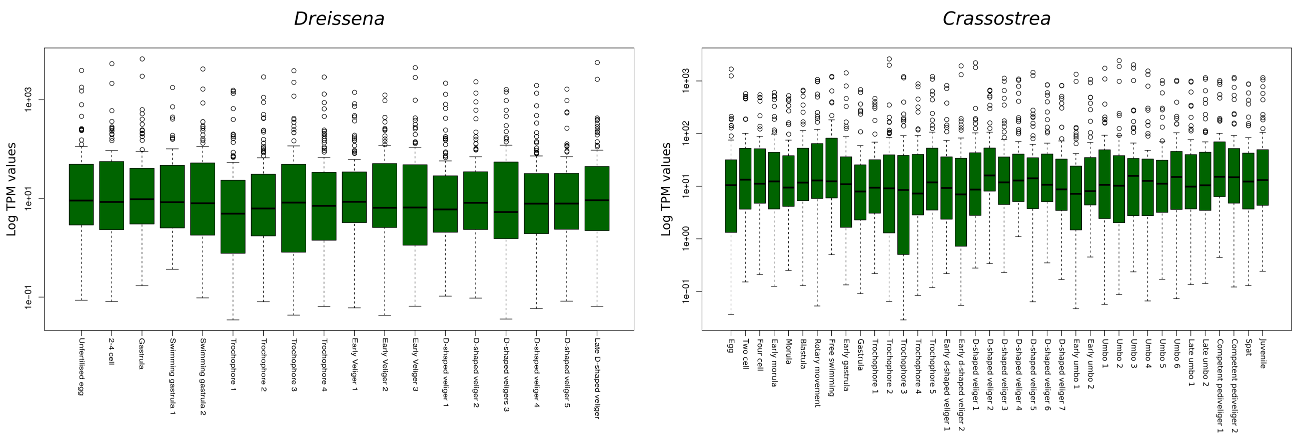

**Figure S15** - Boxplots of the candidate osmoregulation, ionic homeostasis and excretion genes over the course of development in *Dreissena* and *Crassostrea*.

The TPM output of Kallisto was used to perform a Fisher's exact test in R (*40*), comparing the expression levels of the target genes in the early non-swimming developmental stages (unfertilised eggs, 2-4 cell embryos, gastrulas) to background, defined as the average TPM value of all remaining developmental stages. Each was normalised with the scale function in R. An e-value cutoff for significant upregulation was defined as 1e-6 and in total, one aquaporin (Gene.75921), a NHE (Gene.62031) a vacuolar ATPase subunit a (Gene.62284) and a sodium potassium ATPase (Gene.85204) were found to be significantly upregulated in early *Dreissena* development (Fig. S16). This test was adapted from that used for Pfam family expansion by Albertin et al. (2015).

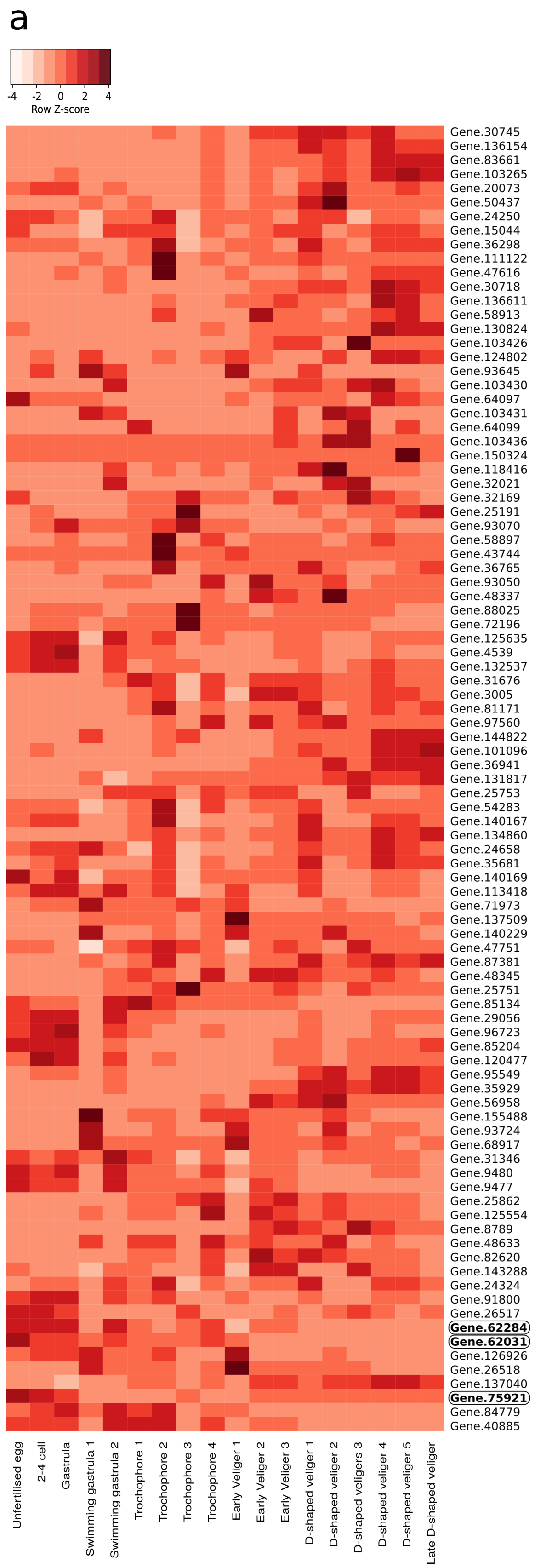

**Figure S16 -** Heatmap of candidate osmoregulatory genes during quagga mussel development. Genes significantly upregulated in early *Dreissena* development but not upregulated during equivalent stages in *Crassostrea* are indicated in bold.

To compare these results to that of a marine species, the same tests were performed with a set of developmental RNA-seq libraries for *Crassostrea gigas* (SRA Bioproject: PRJNA146329). HMMERSEARCH identified 15 aquaporins (Pfam: PF00230.19), two sodium/potassium ATPases (Pfam: PF00287.17), 11 NHEs (Pfam: PF00999.20), eight hydrogen/carbonate co-transporters (Pfam: PF00955.20), 6 voltage gated chloride channels (PF00654.19), 10 cation ATPases (Pfam: PF00689.20, PF13246.5, PF00690.25) and 11 hydrogen ATPases (Pfam: PF00006.24, PF02874.22, PF01813.16, PF03223.14, PF03179.14, PF01496.18, PF01992.15, PF01991.17) from the *Crassostrea* Ensemble (*41*) v9 transcriptome. Using the same settings as for *Dreissena*, we identified one hydrogen/carbonate cotransporter (EKC18553), two sodium/potassium ATPases (EKC41758, EKC32470) and one cation ATPase (EKC34610) that were significantly upregulated in early *Crassostrea* development (Figs. S17, S18).

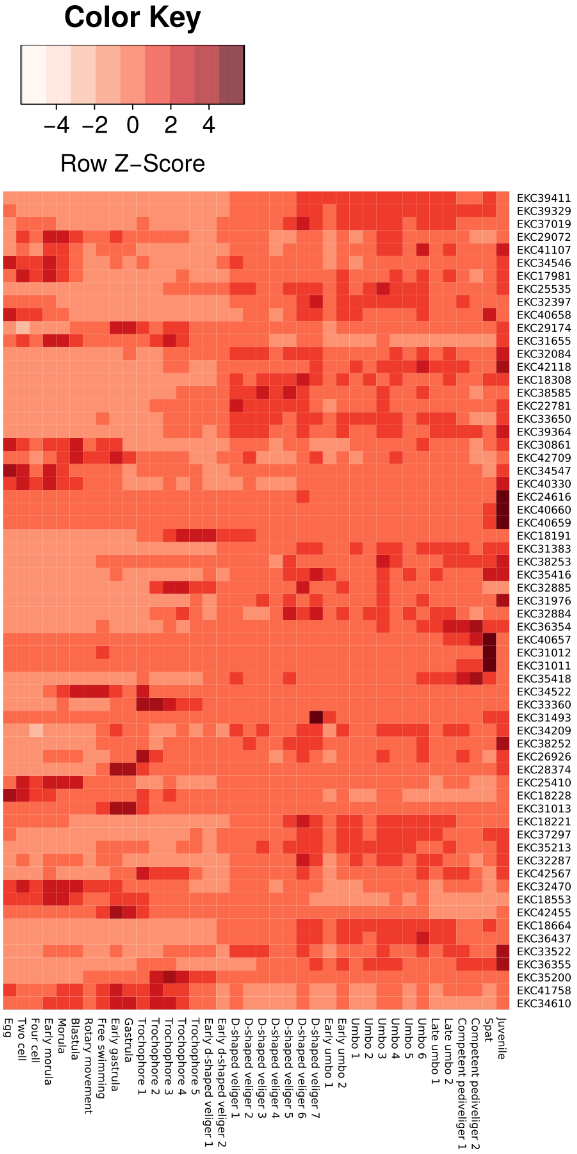

**Figure S17 -** Heatmap of expression of candidate osmoregulatory genes in *Crassostrea gigas* during development.

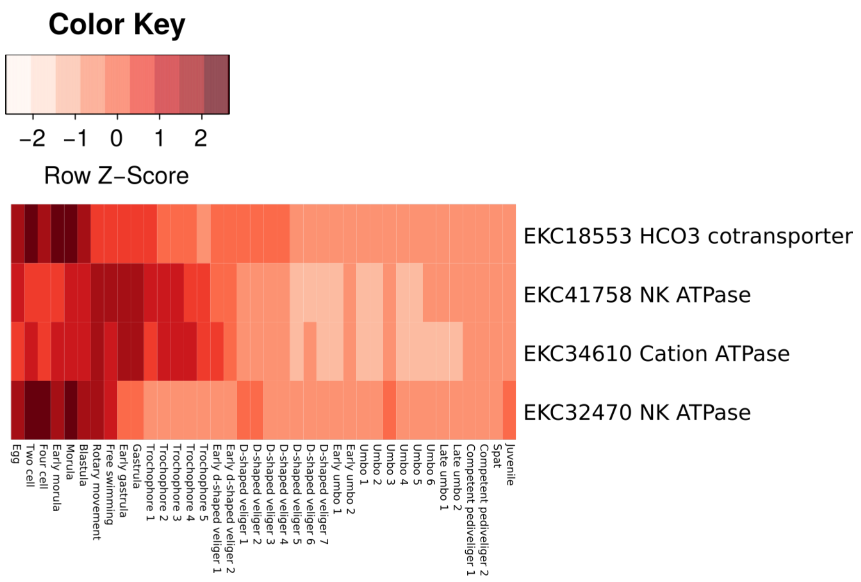

**Figure S18 -** Heatmap of significantly upregulated osmoregulatory genes in early *Crassostrea* development. Early development covers the non-swimming stages (egg, two-cell, four-cell, early morula, morula and blastula).

As both *Dreissena* and *Crassostrea* possess sodium/potassium ATPases that are highly expressed early in development, it was decided not to investigate the role of this gene in *Dreissena* in relation to osmoregulation any further. A direct comparison of the unweighted aquaporin expression levels in *Dreissena* and *Crassostrea* reveals that the most highly expressed aquaporin in the quagga mussel is roughly an order of magnitude more abundant than the most highly expressed aquaporin in the oyster (Fig. S19).

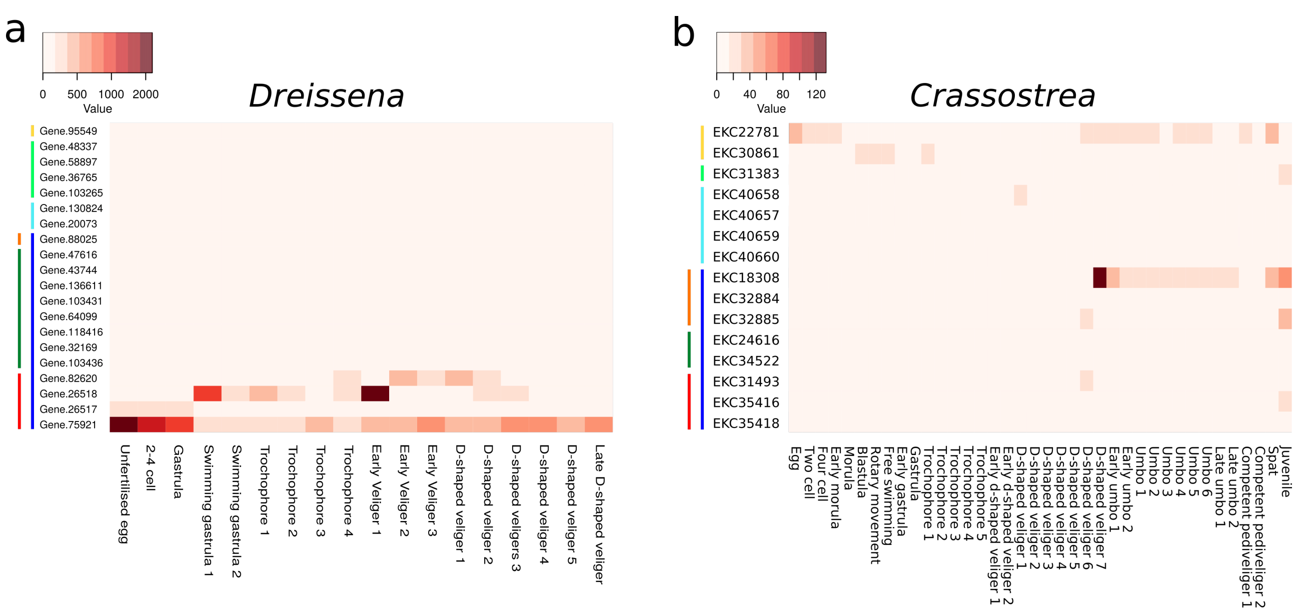

**Figure S19 -** Unweighted expression levels (TPM) of the aquaporin complement of *Dreissena* and *Crassostrea* over development. Of note is the difference in scale between the species. Coloured lines to the left of each graph indicate the class of aquaporin: yellow - unorthodox, light green - aquaglyceroporins, light-blue - aquaammoniaporins, blue - classical aquaporins. The classical aquaporins are further divided: orange - malacoaquaporins, dark green - aqp4-like, red - lophotrochoaquaporins (See SM 7.2).

### SM 7 - Phylogenetics of candidate osmoregulatory genes

#### SM 7.1 - Curation of transcriptomic datasets

To identify orthologues of the candidate osmoregulatory genes in other species, transcriptomes were either downloaded or constructed from publicly available datasets. For those species for which are well curated genome-based transcriptome was available (Annelids: *Helobdella robusta* - JGI (*42*), Bivalves: *Bathymodiolus platifrons* - Dryad (*43*), *Crassostrea gigas* - EnsemblMetazoa (*44*), *Limnoperna fortunei* - GigaDB (*45*), *Modiolus philippinarum* - Dryad (*43*), *Patinopecten yessoensis* - PyBase (*20*), *Pinctada fucata* - OIST marine genomics database (*46*), Cephalochordates: *Branchiostoma floridae* - JGI (*47*), Cephalopods: *Octopus bimaculoides* - UniProt (*48*)*,* Chordates: *Rattus norvegicus* - Ensembl (*49*), Cnidarians: *Nematostella vectensis* - UniProt (*50*), Gastropods: *Biomphalaria glabrata* - UniProt (*51*), *Lottia gigantea* - UniProt (*42*), Echinoderms: S*trongylocentrotus purpuratus* - UniProt (*52*), Insects: *Drosophila melanogaster* - FlyBase (*53*)), these were downloaded from their respective locations. For those species which have not had their genome sequenced, RNA-seq datasets were downloaded from the SRA database (Table S10).

**Table S10** - RNA-seq datasets used for transcriptome construction

| Species | Class | Library number | Accession |
|---|---|---|---|
| Corbicula fluminea | Bivalvia | 1 | SRR5512046 |
| Dreissena polymorpha | Bivalvia | 1 | SRR5000302 |
| Elliptio complanata | Bivalvia | 1 | SRR5136461 |
| Elliptio complanata | Bivalvia | 2 | SRR5136462 |
| Elliptio complanata | Bivalvia | 3 | SRR5136463 |
| Elliptio complanata | Bivalvia | 4 | SRR5136464 |
| Elliptio complanata | Bivalvia | 5 | SRR5136465 |
| Elliptio complanata | Bivalvia | 6 | SRR5136466 |
| Elliptio complanata | Bivalvia | 7 | SRR5136467 |
| Elliptio complanata | Bivalvia | 8 | SRR5136468 |
| Lampsilis cardium | Bivalvia | 1 | SRR1560282 |
| Margaratifera Margaratifera | Bivalvia | 1 | SRR5230914 |
| Mya arenaria | Bivalvia | 1 | SRR1560361 |
| Neotrigonia margaritacea | Bivalvia | 1 | SRR1560432 |
| Villosa lienosa | Bivalvia | 1 | SRR354206 |
| Villosa lienosa | Bivalvia | 2 | SRR354207 |
| Lymnaea stagnalis | Gastropoda | 1 | SRR6832921 |
| Lymnaea stagnalis | Gastropoda | 2 | SRR6832922 |
| Lymnaea stagnalis | Gastropoda | 3 | SRR6832924 |

RNASeq libraries were processed with Trimmomatic v0.35 (*13*) and assembled with Binpacker (*24*) using the kmer values k23, k25, k27, k32. Individual kmer assemblies were merged with Velvet (*25*) then de-duplicated with cd-hit (*54*) allowing for up to 98% similarity. Open reading frames (ORFs) were predicted with Transdecoder v 3.0.0 using the --single_best_only option (*27*). The only exception to this workflow were the libraries of *E. complanata, D. polymorpha* and *L. stagnalis* where IDBA-tran (*55*) was used for transcriptome construction. For species with more than one available RNA-seq library, separate transcriptomes were produced.

**SM 7.2 - Aquaporin phylogenetics**

The PFam hidden markov model (hmm) for major intrinsic protein (MIP/aquaporin: PF00230) was used to search all species under investigation for candidate aquaporin genes with hmmsearch from the HMMER package v3.1 (*9*). Candidate aquaporins were aligned with MAFFT v7.310 (*8*) and viewed with Aliview v1.21 (*56*). Truncated and duplicated sequences were manually removed from the list of candidates and BMGE v1.12 (*57*) was used to trim the final alignment. Phylogenetic tree construction was conducted with FastTree v2.1.10 (*11*).

The resulting tree successfully resolved the four major animal aquaporin classes - classical aquaporins, aquaamoniaporins, unorthodox aquaporins and aquaglyceroporins (Fig. S20; *58*). We also identified several smaller classes and subclasses including entomoglyceroporins (EGLPs) (*59*), malacoaquaporins (*60*), *Drosophila* intrinsic proteins (DRIPs) (*61*) and *Pyrocoelia rufa* integral proteins (PRIPs) (*62*).

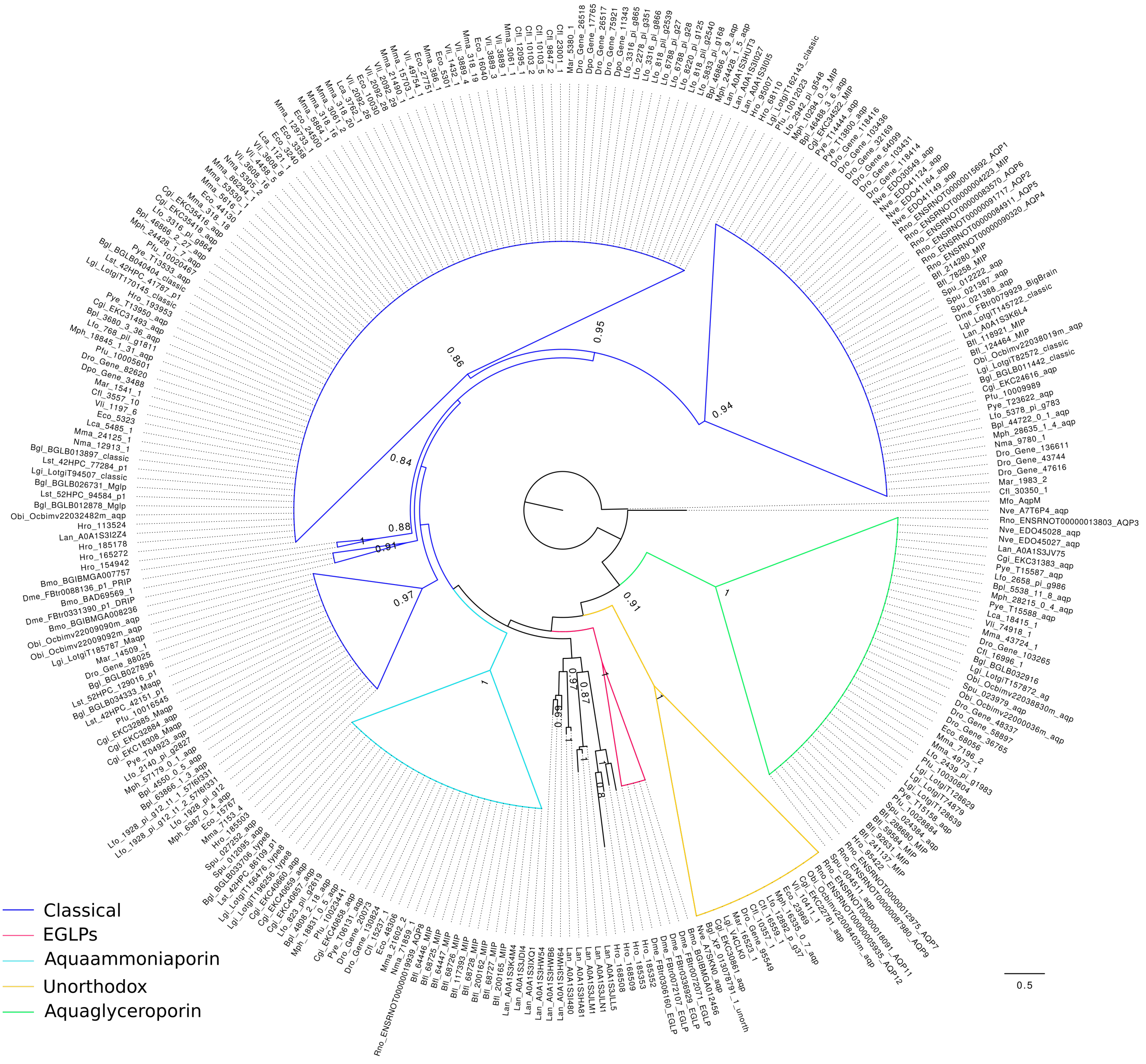

**Figure S20 -** Aquaporin phylogeny. Phylogenetic tree of aquaporins in *Branchiostoma floridae* (Bfl), *Biomphalaria glabrata* (Bgl), *Bombyx mori* (Bmo), *Bathymodiolus platifrons* (Bpl), *Corbicula fluminea* (Cfl), *Crassostrea gigas* (Cgi), *Drosophila melanogaster* (Dme), *Dreissena polymorpha* (Dpo), *Dreissena rostriformis* (Dro), *Elliptio complanata* (Eco), *Helobdella robusta* (Hro), *Lingula anatina* (Lan), *Lampsilis cardium* (Lca), *Limnoperna fortunei* (Lfo), *Lottia gigantea* (Lgi), *Lymnaea stagnalis* (Lst), *Mya arenaria* (Mar), *Methanobacterium formicicum* (Mfo), *Margaritifera margaritifera* (Mma), *Modiolus philippinarum* (Mph), *Neotrigonia margaritacea* (Nma), *Nematostella vectensis* (Nve), *Octopus bimaculoides* (Obi), *Pinctada fucata* (Pfu), *Patinopecten yessoensis*

(Pye), *Rattus norvegicus* (Rno), *Strongylocentrotus purpuratus* (Spu), *Villosa lienosa* (Vli). Colours depict aquaporin class as indicated. The classical aquaporins are further divided (left to right) in to malacoaquaporins, DRIPs, PRIPs, lophotrochoaquaporins and aqp4-like. Support values below 0.8 are not displayed.

While the phylogenetic positions of the DRIPs and PRIPs are consistent with previous results, the EGLPs were positioned alongside the aquaamoniaporins and aquaglyceroporins, rather than with the aqp4-like classical aquaporins (Fig. S20; *56*). EGLPs, like aquaaminoporins, aquaglyceroporins and unorthodox aquaporins, are capable of transporting a range of solutes in addition to water (*59*, *63–67*). A closer relationship between these groups to the exclusion of the classical aquaporins may reflect an ancestral character state. In addition to the major aquaporin clades, a group consisting of representatives from *Lingula* and *Helobdella* were located between the aquaammoniaporins and the EGLPs. The function and position of this clade will require further confirmation. No support for the malacoglyceroporin (Mglp) clade was found in our study (*60*, *68*). As no evidence exists for the transport of glycerol by these proteins and due to their phylogenetic position amongst the classical aquaporins, we suggest discontinuation of the term 'malacoglyceroporin'.

The largest group of aquaporins identified in our analysis is a previously unreported clade of classical aquaporins consisting solely of lophotrochozoan representatives (Fig. S20). We refer to this clade as the lophotrochoaquaporins. In the quagga mussel, three lophotrochoaquaporin orthologues form a clade with those of the congeneric freshwater *Dreissena polymorpha* (two orthologues) and the closely related marine species *Mya arenaria* (one orthologue). While the five orthologues of the freshwater *Corbicula fluminea* were not annotated against a genome, the level of sequence divergence between the orthologues makes it likely that at least three represent true paralogues (Fig. S21). This is the same number of paralogues present in the quagga mussel.

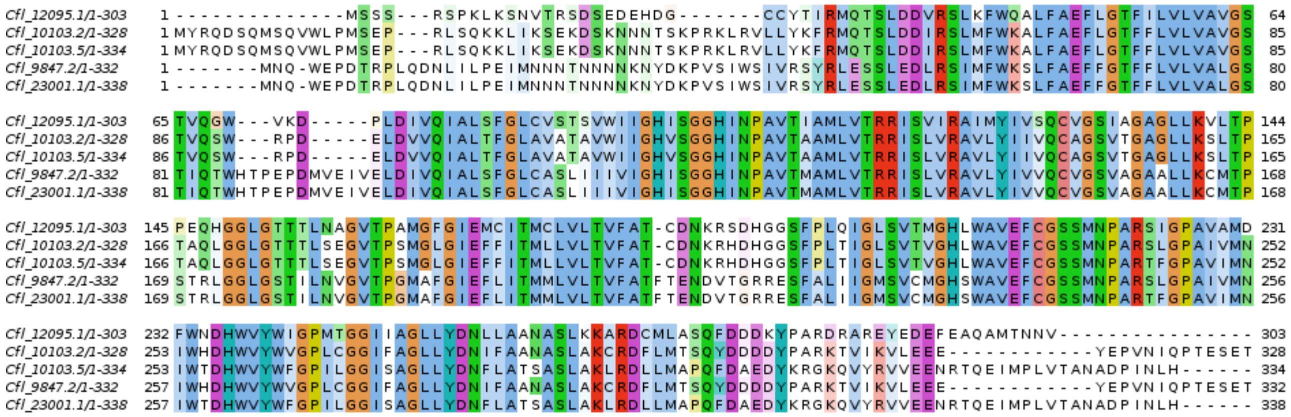

**Figure S21** - Multiple sequence alignment of *Corbicula* lophotrochoaquaporins aquaporins. The five copies identified may represent three paralogues in addition to isoforms of two of the gene copies. Definitive determination of this possibility will require annotation with a genome assembly.

The distantly related freshwater golden mussel *Limnoperna fortunei* possess an expanded set of nine lophotrochoaquaporin orthologues whereas its closest relatives, the marine *M. philippinarum* and *B. platifrons*, possess only a single copy each. No genomic resources are yet available for any of the freshwater paleoheterodonts (unionids) and so to avoid conflating transcript variants with true orthologues, expanded aquaporin clades were only annotated if they included representatives from at least three species. Four unionid clades were identified, indicating that the last common ancestor of these species likely already possessed an expanded repertoire of lophotrochoaquaporins. Only a single lophotrochoaquaporin orthologue was identified in the marine paleoheterodont *N. margaritacea* (Fig. S20). While the identification of more *N. margaritacea* orthologues with increased sampling cannot be ruled out, it appears that the paleoheterodont lophotrochoaquaporin expansion occurred after the divergence of the marine and freshwater species and before speciation of the unionids.

Outside of the molluscs, the freshwater annelid leech *H. robusta* also appears to have an expanded set of lophotrochoaquaporins (three orthologues). No such expansion was identified in the freshwater gastropods *B. glabrata* or *L. stagnalis*.

**SM 7.3 - v-ATPase subunit a phylogentics**

The v-ATPase subunit a is the most diverse of the v-ATPase subunits (*69*, *70*) and is responsible for targeting the v-ATPase complex to specific sites within the cell (*71*, *72*). The PFam hmm (PF01496.18) was used to search for v-ATPase subunit a as per the aquaporin orthologues (SM 7.2). Alignment, processing and phylogenetic tree construction were also conducted as detailed in SM 7.2. In total 43 orthologues comprised the final alignment.

No comprehensive phylogeny of metazoan v-ATPase subunit a sequences is yet available. In vertebrates, four subunit a isoforms have been identified, each with distinct functions and expression patterns (*73*, *74*). It is unknown how these relate to the v-ATPase subunit a isoforms of other metazoan lineages. Our analyses reveals v-ATPase subunit a expansions in each phylum investigated, however molluscs appear to have undergone two rounds of diversification giving rise to two distinct monophyletic subclasses (Fig. S22). The quagga mussel v-ATPase subunit a orthologue found to be highly expressed during early embryogenesis (*Gene.62284*) is a member of subclass II however no consistent pattern of expansion in freshwater bivalves akin to that of aquaporins was observed.

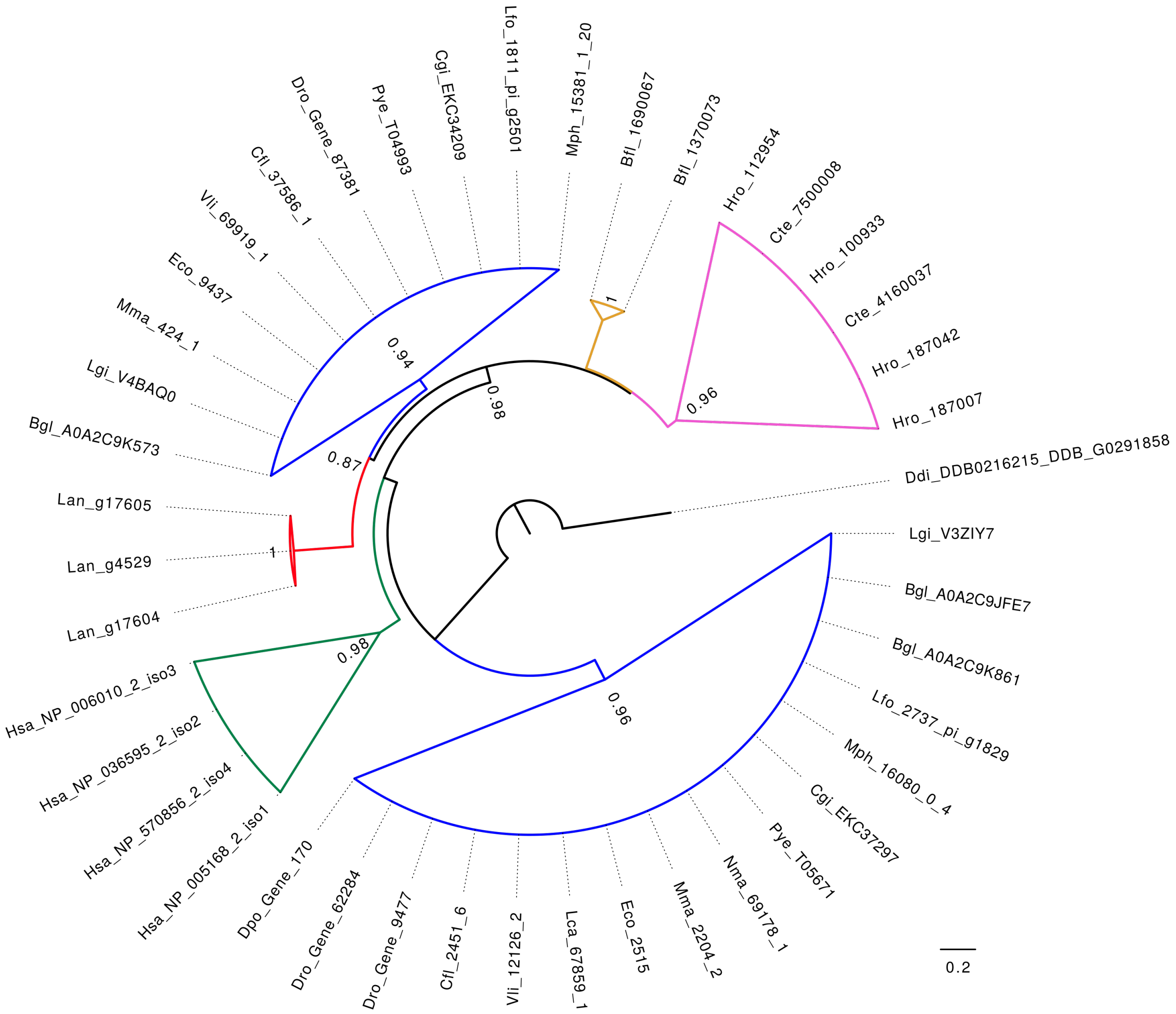

**Figure S22 -** Vacuolar ATPases. Phylogenetic tree of vATPases in *Branchiostoma floridae* (Bfl), *Biomphalaria glabrata* (Bgl), *Corbicula fluminea* (Cfl), *Crassostrea gigas* (Cgi), *Capitella teleta* (Cte), *Dictyostelium discoideum* (Ddi), *Dreissena polymorpha* (Dpo), *Dreissena rostriformis* (Dro), *Elliptio complanata* (Eco), *Helobdella robusta* (Hro), *Homo sapiens* (Hsa), *Lingula anatina* (Lan),

*Lampsilis cardium* (Lca), *Limnoperna fortunei* (Lfo), *Lottia gigantea* (Lgi), *Margaritifera margaritifera* (Mma), *Modiolus philippinarum* (Mph), *Neotrigonia margaritacea* (Nma), *Patinopecten yessoensis* (Pye), *Villosa lienosa* (Vli). Depicted are phyletic expansions of vATPases in vertebrates (green), brachiopods (red), cephalochordates (orange), annelids (pink) and two in molluscs (blue), each of which is collapsed in the figure. Support values under 0.8 are not displayed.

#### SM 7.4 - Sodium hydrogen exchanger phylogenetics

Sodium hydrogen exchangers form part of the monovalent cation proton antiporter (CAP) superfamily (*75*). CAP orthologue identification (Pfam: PF00999.20), alignment, processing and phylogenetic tree construction proceeded as per SM 7.2. The resulting tree successfully resolved all previously reported animal CAP classes - NHA, PM-NHE and Endo/TGN IC-NHE and NHE8-like IC-NHE (Fig. S23). We were also able to resolve the position of the enigmatic mammalian sperm NHEs with a well supported clade consisting of deuterostome, lophotrochozoan and ecdysozoan orthologues, in addition to the plant SOS1 sequences. The non-animal clades CHX, NhaP and plant vacuolar were also resolved. We also found support for two previously unreported animal CAP clades. The first, consisting of deuterostome and lophotrochozoan sequences, is most closely aligned to the plant CHX transporters. No molluscan sequences were identified from this clade. The second was a large lophotrochozoan-specific family of NHEs most closely aligned to the PM-NHEs found in all major animal superphyla.

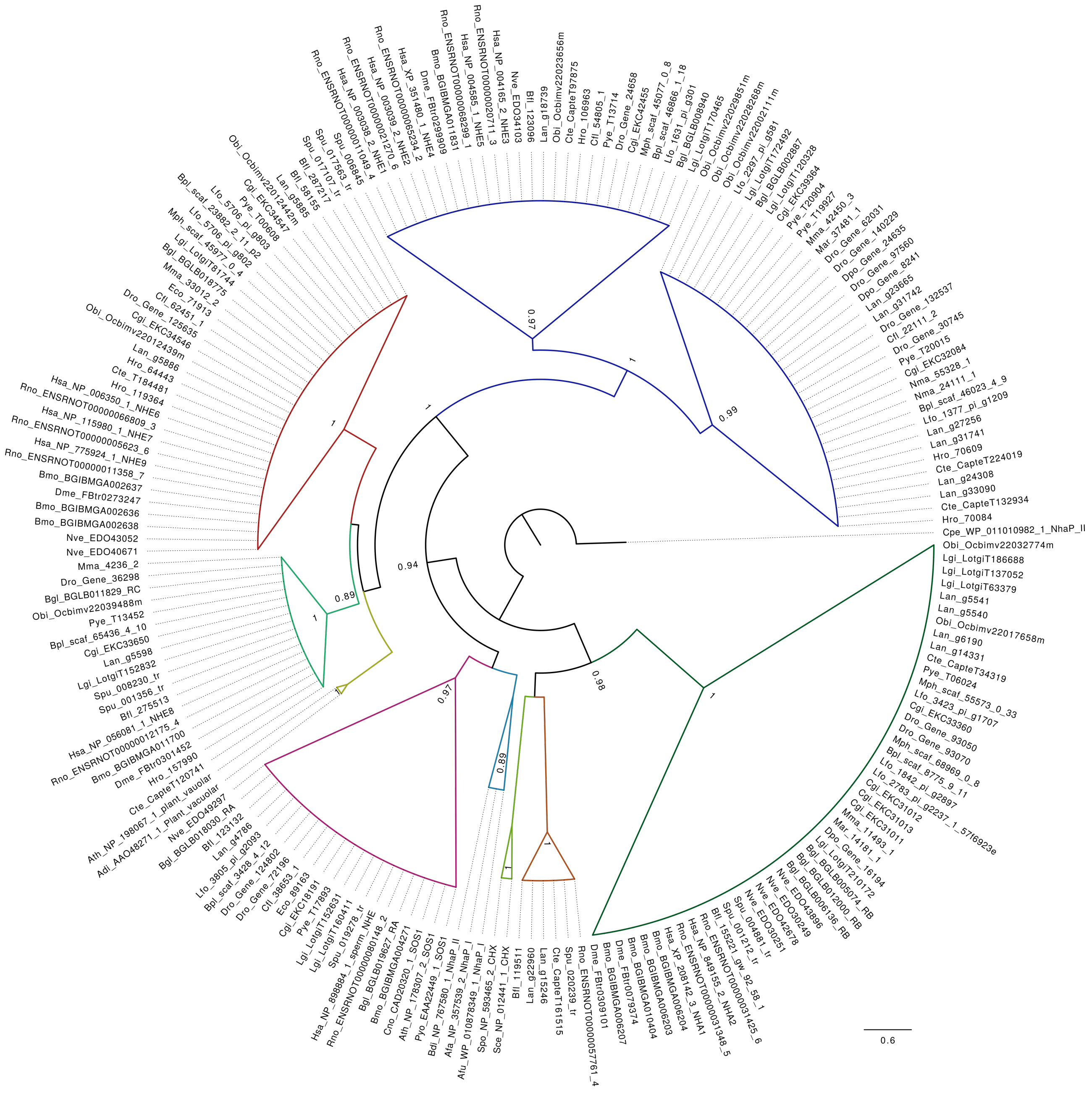

**Figure S23 -** Monovalent cation proton antiporter (CAP) families. Phylogenetic tree of CAPs in *Atriplex dimorphostegia* (Adi), *Agrobacterium fabrum* (Afa), *Archaeoglobus fulgidus* (Afu), *Arabidopsis thaliana* (Ath), *Bradyrhizobium diazoefficiens* (Bdi), *Branchiostoma floridae* (Bfl), *Biomphalaria glabrata* (Bgl), *Bombyx mori* (Bmo), *Bathymodiolus platifrons* (Bpl), *Corbicula fluminea* (Cfl), *Crassostrea gigas* (Cgi), *Cymodocea nodosa* (Cno), *Clostridium perfringens* (Cpe), *Capitella teleta* (Cte), *Drosophila melanogaster* (Dme), *Dreissena polymorpha* (Dpo), *Dreissena rostriformis* (Dro), *Elliptio complanata* (Eco), *Helobdella robusta* (Hro), *Homo sapiens* (Hsa), *Lingula anatina* (Lan), *Limnoperna fortunei* (Lfo), *Lottia gigantea* (Lgi), *Mya arenaria* (Mar), *Margaritifera margaritifera* (Mma), *Modiolus philippinarum* (Mph), *Neotrigonia margaritacea* (Nma), *Nematostella vectensis* (Nve), *Octopus bimaculoides* (Obi), *Patinopecten yessoensis* (Pye), *Plasmodium yoelii* (Pyo), *Rattus norvegicus* (Rno), *Saccharomyces cerevisiae* (Sce),

*Schizosaccharomyces pombe* (Spo), *Strongylocentrotus purpuratus* (Spu) with each faimly collapsed. From the top of the image and moving clockwise, plasma membrane (PM)-NHEs including lophotrochozoan-specific clade (blue), sodium-hydrogen antiporter (NHA, dark green), undescribed CHX-like clade (orange), CHX (green), NhaP (light blue), SOS1/mammalian sperm (fuschia), plant vacuolar (yellow), intracellular (IC) NHE8-like (lime), intracellular (IC) Endo/TGN (red).

The quagga mussel NHE found to be highly expressed during early embryogenesis (*Gene.62031*) is a member of this lophotrochozoan-specific NHE family. While five quagga mussel orthologues were identified in this clade, as with the v-ATPase subunit a, no pattern of expansion in freshwater bivalves was observed.

### SM 8 - Lophotrochoaquaporin structural modelling

#### SM 8.1 - 3D structural modelling of Dro.75921

The highly expressed *Dreissena* lophotrochoaquaporin orthologue (*Dro.75921*) sequence was uploaded to SWISS-MODEL (*76*) for structural modelling. Models were built from the top 14 templates (as determined by a quaternary structure quality estimate (QSQE) of greater than 0.5), which included structures from four aquaporin orthologues - *AQPO* (PDB: 1YMG), *AQP1* (PDB: 5C5X), *AQP4* (PDB: 1J4N) and *AQP5* (PDB: 2ZZ9). For all four orthologues, the most structurally variable regions as measured by the QMEAN score corresponded to loops A, C and D with the exception of *AQP1* which showed strong structural similarity to *Dro.75921* through loop D with the QMEAN not dropping below 0.68 (SWISS-MODEL cut off for low quality equals 0.6) (Fig. S24). Lens fibre major intrinsic protein, *AQP4* and *AQP5* have minimum QMEAN scores through loop D of 0.48, 0.48 and 0.58 respectively.

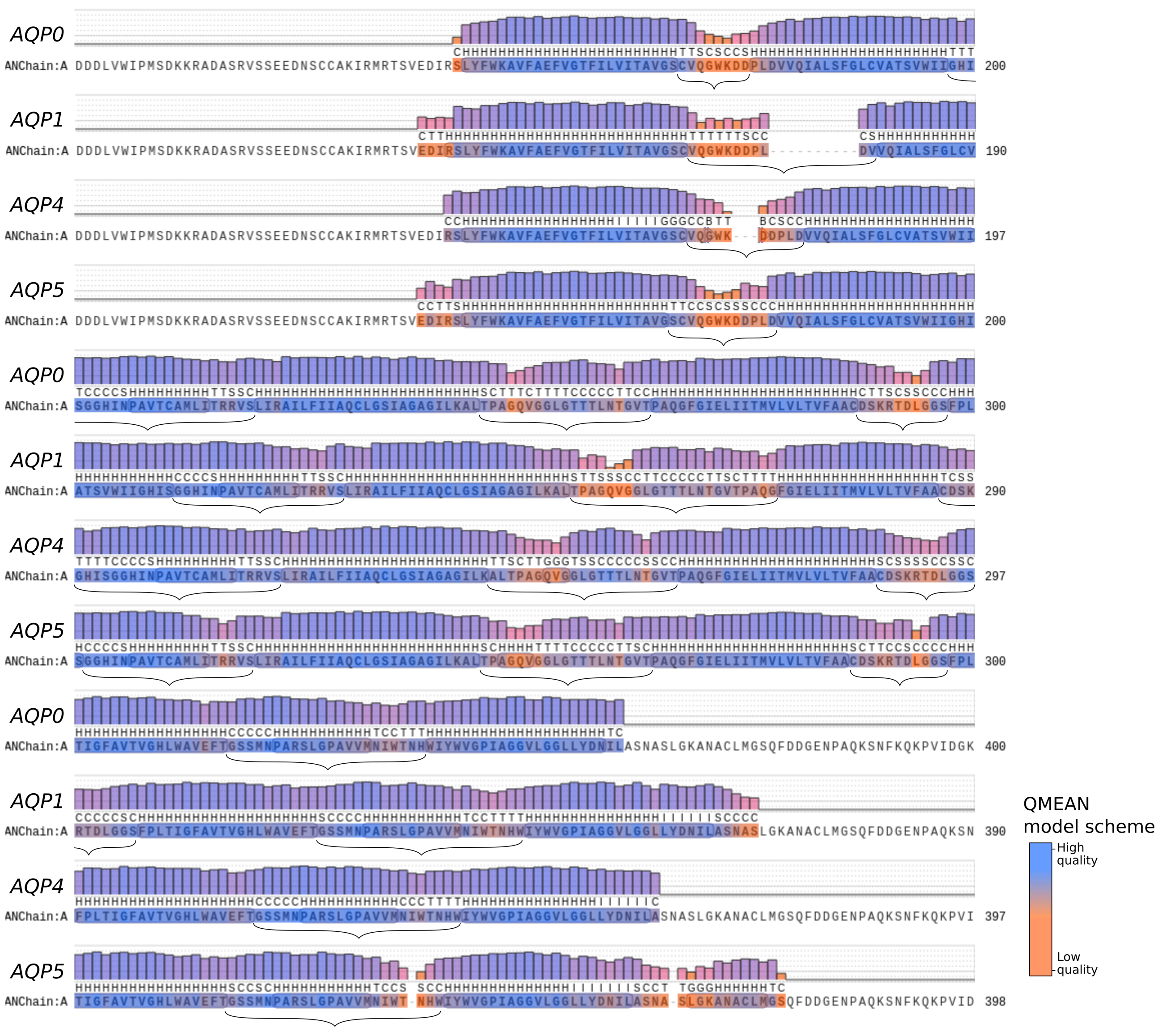

**Figure S24 -** Model-template alignment of *Dro.7592*1 wrapped to *AQP0* (PDB: 1ymg.1), *AQP1* (PDB: 1j4n.1), *AQP4* (PDB: 1j4n.1) and *AQP5* (PDB: 5c5x.1). Each position is colour-coded according to the QMEAN model quality score and loop regions are indicated with brackets (loops A-E). Of note is loop D (fourth loop in each sequence) which shows structural similarity with the loop D of *AQP1* but low structural similarity with the other three models.

Loop D has been implicated in the gating of aquaporins and is hypothesised to impact the flux of both water and ions (*77–80*). As can be seen in Fig. S25, loop D forms a point of restriction on the central tetrameric pore and being composed of highly charged residues, is likely to impact the flux of charged particles.

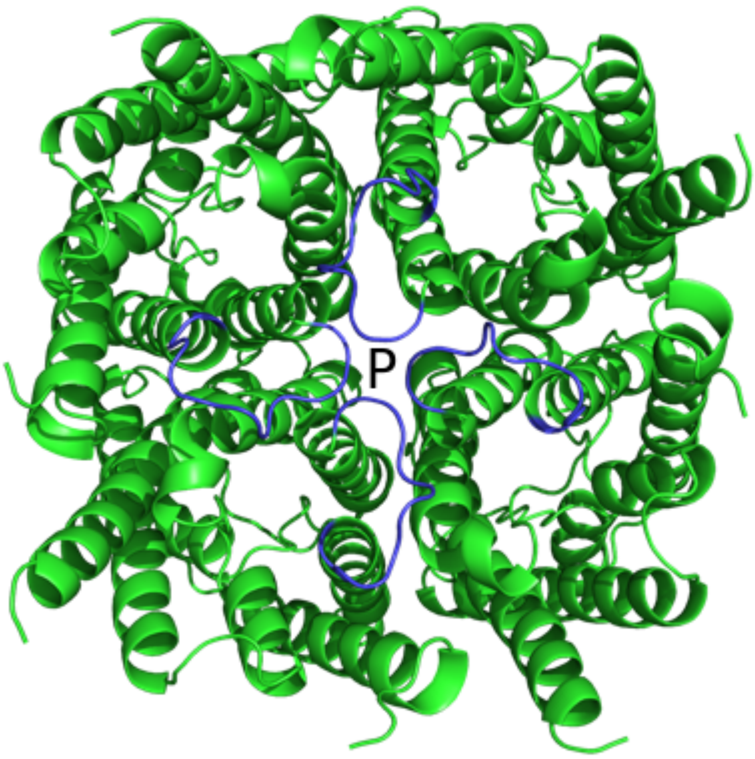

**Figure S25 -** Cytoplasmic view of the predicted quaternary structure of tetrameric *Dro.75921* wrapped to *AQP1* (PDB: 1j4n.1) with the loop D of each chain coloured blue and the central tetrameric pore indicated (P).

**SM 8.2 - Classical aquaporin loop D structure and conservation**

Stabilisation of the quaternary structure through the formation of salt bridges was predicted with ESBRI (*81*) revealing a bridge between Arg-218 in the conserved NPA-motif containing loop B and Asp358 located in helix VI, and a bridge between Arg-291 and Asp-293 both located within loop D (Table S11; Fig. 2). Salt bridges are non-covalent bonds between oppositely charged residues that are located in close physical proximity to one another within a folded protein. As such, the strength of the interaction is subject to physiological pH and large changes to pH, both high and low, can impact the protonation of charged amino acid residues (*82*, *83*).

**Table S11 -** Predicted salt bridges in *Dro.75921*

| Residue 1 | Residue 2 | Distance |
|---|---|---|
| NH1 ARG A 218 | OD1 ASP A 358 | 3.11 |
| NH2 ARG A 291 | OD1 ASP A 293 | 2.67 |
| NH2 ARG A 291 | OD2 ASP A 293 | 3.37 |
| NH1 ARG B 218 | OD1 ASP B 358 | 3.11 |
| NH2 ARG B 291 | OD1 ASP B 293 | 2.67 |
| NH2 ARG B 291 | OD2 ASP B 293 | 3.37 |
| NH1 ARG C 218 | OD1 ASP C 358 | 3.11 |
| NH2 ARG C 291 | OD1 ASP C 293 | 2.67 |
| NH2 ARG C 291 | OD2 ASP C 293 | 3.37 |
| NH1 ARG D 218 | OD1 ASP D 358 | 3.11 |
| NH2 ARG D 291 | OD1 ASP D 293 | 2.67 |
| NH2 ARG D 291 | OD2 ASP D 293 | 3.37 |

To identify differences in the loop D of aqp4-like aquaporins and lophotrochoaquaporins, peptide logos (*84*) for each were produced from alignments corresponding to the loop D region. Logos were made for the aqp4-like sequences, the lophotrochozoan representatives of the aqp4-like aquaporins only and for the lophotrochoaquaporins sequences (Fig. S26).

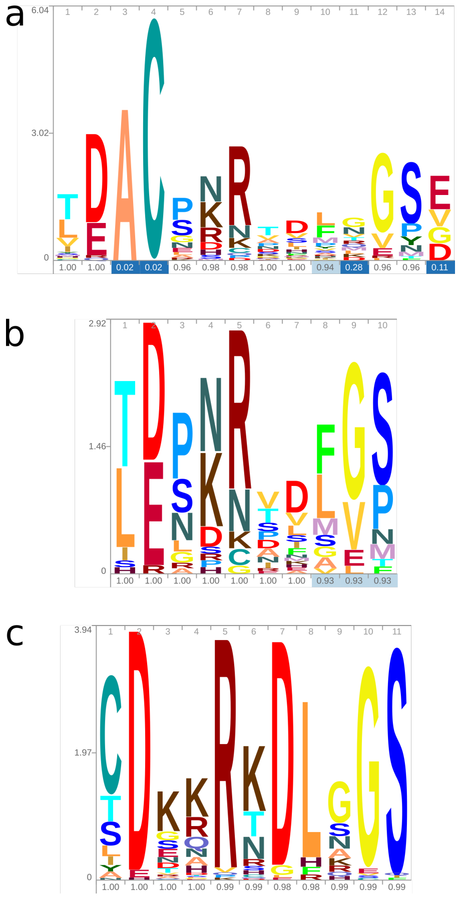

**Figure S26 -** Peptide logos for aquaporin loop D regions. a) Logo for all aqp4-like sequences (49 sequences), b) Logo for lophotrochozoan representatives of aqp4-like aquaporins (29 sequences), c) Logo for lophotrochoaquaporin sequences (98 sequences).

This revealed considerable conservation amongst the lophotrochoaquaporins and more variability amongst the aqp4-like sequences. In particular, the two residues predicted to form a salt bridge in loop D (Arg-291 and Asp-293) are two of the most highly conserved in lophotrochoaquaporins.

**SM 9 - Embryogenesis and its response to osmolarity challenges**

**SM 9.1 - Development and cleavage cavity formation under ambient conditions**

To observe development of the embryos under ambient conditions, adult mussels collected from the Danube river in Vienna, Austria, were induced to spawn through immersion in a serotonin solution as per SM 5.2. *Dreissena* eggs possess a thick jelly coat that prevents them from concentrating to high density when kept in a mono-layer. This prevents the microscopic observation of a large number of eggs at a time as the low density means few eggs are present in the microscope field of view.

To remove the jelly coat, a protocol was developed based on one designed to remove the jelly coat of eggs from the sea urchin *Strongylocentrotus purpuratus* (*85*). Eggs were moved from ambient FRW at pH 8.6 to low pH FRW (pH 5.7) and gently mixed for two minutes, whereafter they were thoroughly washed with ambient FRW. Treated eggs were then transferred to a 50ml tube which was continuously inverted for three minutes before being centrifuged at 500xG for three minutes. FRW was decanted and replaced and the tube inversion and centrifugation was repeated three more times (four in total) producing viable de-jellied eggs (Fig. S27). De-jellied eggs were fertilised through the introduction of sperm for 15 minutes, after which the eggs were washed thoroughly with FRW. Fertilised eggs were transferred to a WillCo glass bottom dish for observation on a Leitz Labovert inverted microscope. Video recordings began 45 minutes post fertilisation and ran for 5.25 hours to capture embryos up to the point of six hours post fertilisation (hpf). This corresponded to the point when most embryos had reached the swimming gastrula stage of development. Ambient conditions meant that development occurred at approximately 26 degrees Celsius.

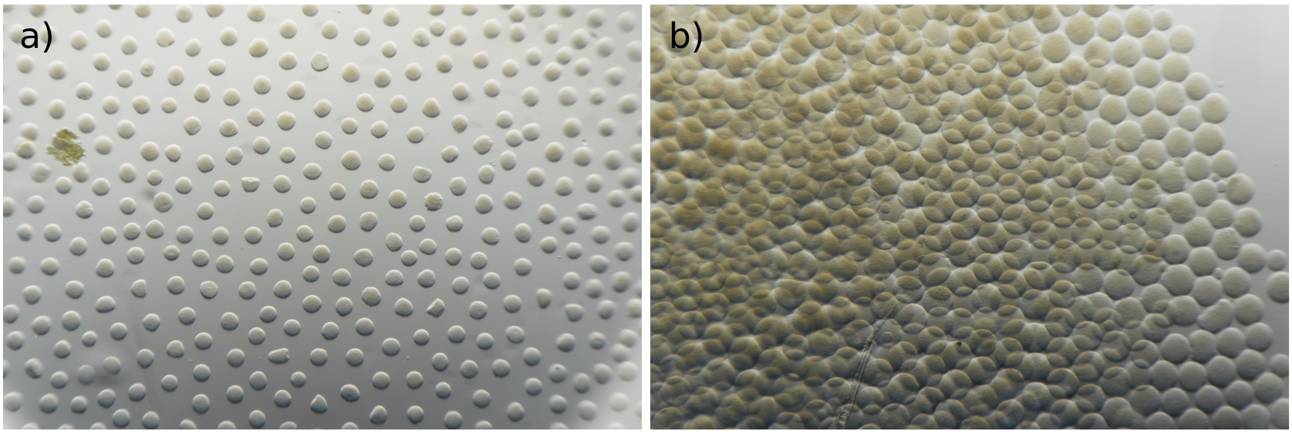

**Figure S27 -** Egg jelly layer removal. a) Untreated eggs form a mono-layer where individual eggs are prevented from touching due to the presence of a thick transparent jelly layer (not visible). b) Treated eggs are no longer prevented from accumulating in high densities due to the absence of the jelly layer.

First cleavage was observed to occur at about 1 hpf in both treated and untreated samples with the second and third occurring at approximately one hour intervals. No impact on fertilisation or development as a consequence of jelly coat removal was observed. The first cleavage cavity began to form between the two daughter cells shortly after the first cleavage was completed. As has been observed in several species, cleavage cavity formation begins simultaneously at several positions along the cell-cell margin resulting in small lens-shaped fluid filled cavities, which gradually grow and coalesce until a single cavity remains (Fig. S28). Cleavage cavities often grow to occupy a substantial proportion of the total embryo volume leaving only a small ring traversing the circumference of the embryo where cell-cell contact remains intact. Eventually this final ring of contact is breached by the growing cavity whereupon the fluid in the cavity is rapidly discharged, possibly through the release of tension built up in the fertilisation envelope. This discharge results in the collapse of the cavity (Fig. S28).

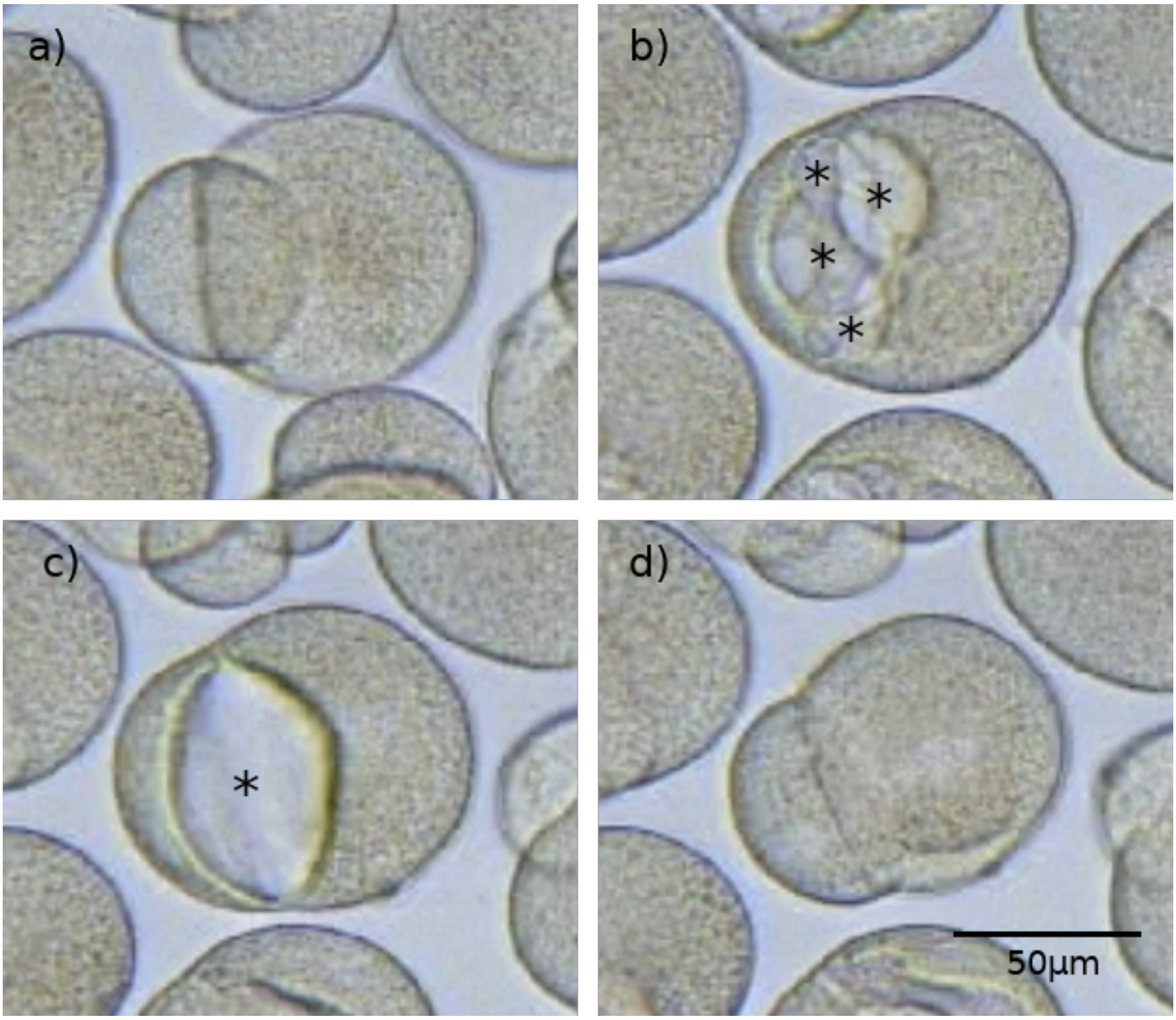

**Figure S28 -** Development under ambient conditions. a) Newly cleaved two-cell embryo. b) Two-cell embryo with numerous small cleavage cavities (*) located along the cell-cell margin. c) Two-cell embryo with a single large cleavage cavity (*) following the coalescing of the numerous small early cavities. d) Two cell embryo immediately following cleavage cavity collapse.

The process of cleavage cavity inflation and collapse typically repeats two to four times during each of the first three cleavages. At the eight-cell stage, a blastocoel forms and this also appears to periodically inflate and collapse in a similar way to the cleavage cavities (Fig. S29).

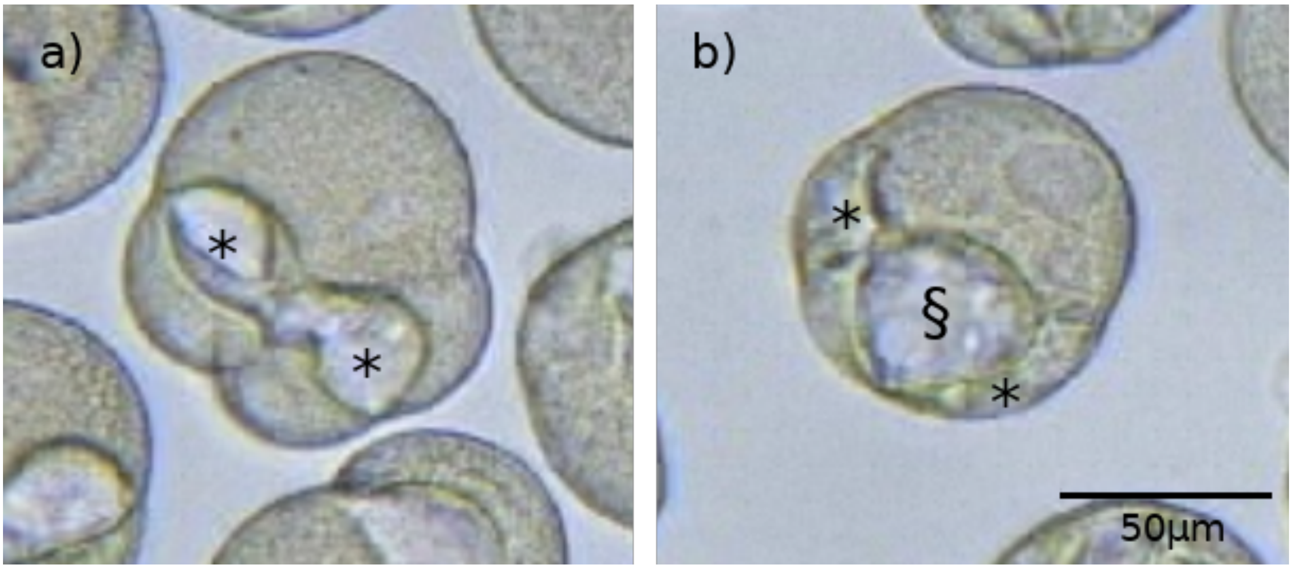

**Figure S29 -** Second and third cleavage. a) Four-cell embryo with prominent cleavage cavities (*). b) Eight-cell embryo with cleavage cavities and inflated blastocoel (§).

Over the six hours of development that were recorded, most embryos remained at roughly the volume at which they began, not withstanding the repeated inflation and collapse attributable to cleavage cavity activity. In contrast, those eggs that failed to become fertilised gradually increased in volume over the recording period (Fig. S30). This is most likely due to the osmotic influx of water across the vitelline envelope and cell membrane that, in the absence of cleavage cavity formation, was not able to be excreted.

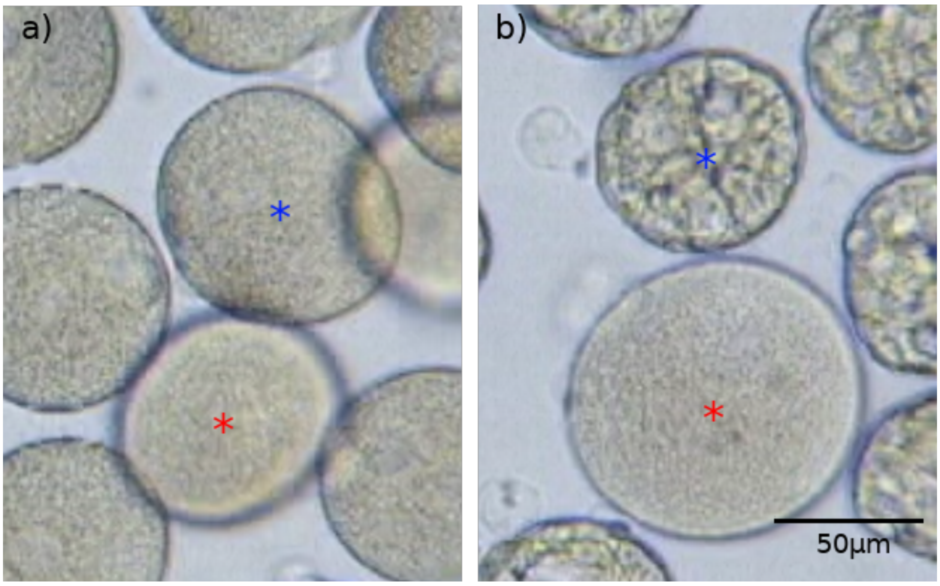

**Figure S30 -** Increase in volume of unfertilised eggs over time. a) Prior to cleavage the zygote (blue star) and the unfertilised egg (red star) are roughly equal in volume. b) Just prior to the swimming gastrula stage, the gastrula (blue star) remains at a roughly equal volume to when it was a zygote while the unfertilised egg (red star) has increased in volume.

On rare occasions under ambient osmotic conditions, cleavage of the embryo fails. We have observed that embryos that are successfully fertilised, as determined by the presence of polar bodies, but fail to cleave correctly are at an increased risk of fertilisation envelope rupture. When the fertilisation envelope ruptures, part of the embryo is extruded from the fertilisation envelope, leading to the formation of large highly active amoeboid processes that often contain large vacuole-like structures (Fig. S31).

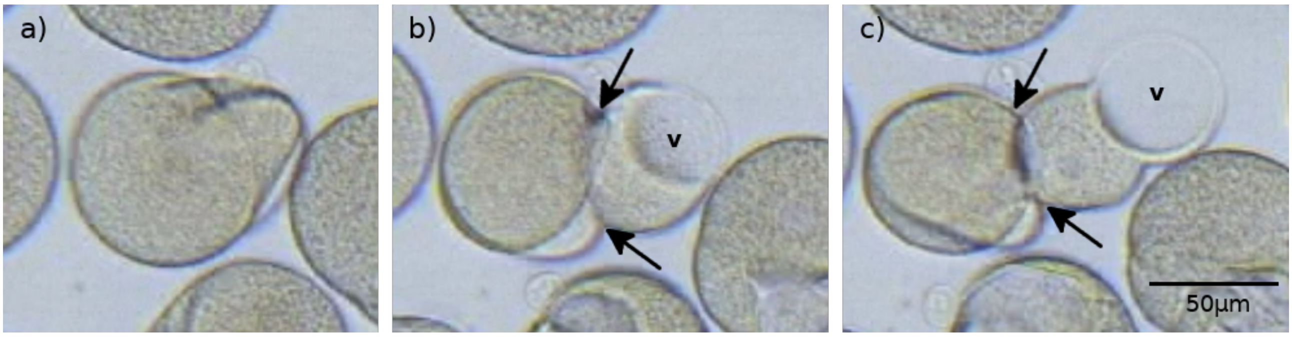

**Figure S31 -** Fertilisation envelope ruptures caused by a failure to cleave. a) Fertilised zygote shortly after an attempted first cleavage with distorted fertilisation envelope. b) Fertilised zygote shortly after rupture of the fertilisation envelope (arrows mark rupture) with a large vacuole-containing (v) process rapidly extruding through the point of rupture toward the right of the zygote. c) Extruded vacuole-containing (v) amoeboid process (right of arrows) of roughly equal volume to the part of the zygote still located within the fertilisation envelope (left of arrows).

**SM 9.2 - Development and cleavage cavity formation under hyperosmotic conditions**

To test the hypothesised function of cleavage cavities as embryonic osmotic regulatory structures, embryos were raised under high salt conditions. Eggs were de-jellied and fertilised in ambient FRW as per SM 8.1 before being transferred to a high salt solution made by dissolving a commercial artificial seawater salt mix (Sera Marin) in FRW. An initial trial tested how development proceeded in five salt concentrations - 3.5 parts per thousand (ppt), 2.63 ppt, 1.75 ppt, 0.875 ppt and 0.35 ppt. In all but the highest concentration, development of most embryos proceeded to at least the gastrula stage and so 1.75 ppt was selected for further experiments.

As in SM 8.1, video recording began 45 minutes post fertilisation. In contrast to the ambient river water, embryos raised in the high salt solution progressed through the cleavage stages either without the production of cleavage cavities, or in a few cases, with highly reduced cleavage cavities (Fig. S32).

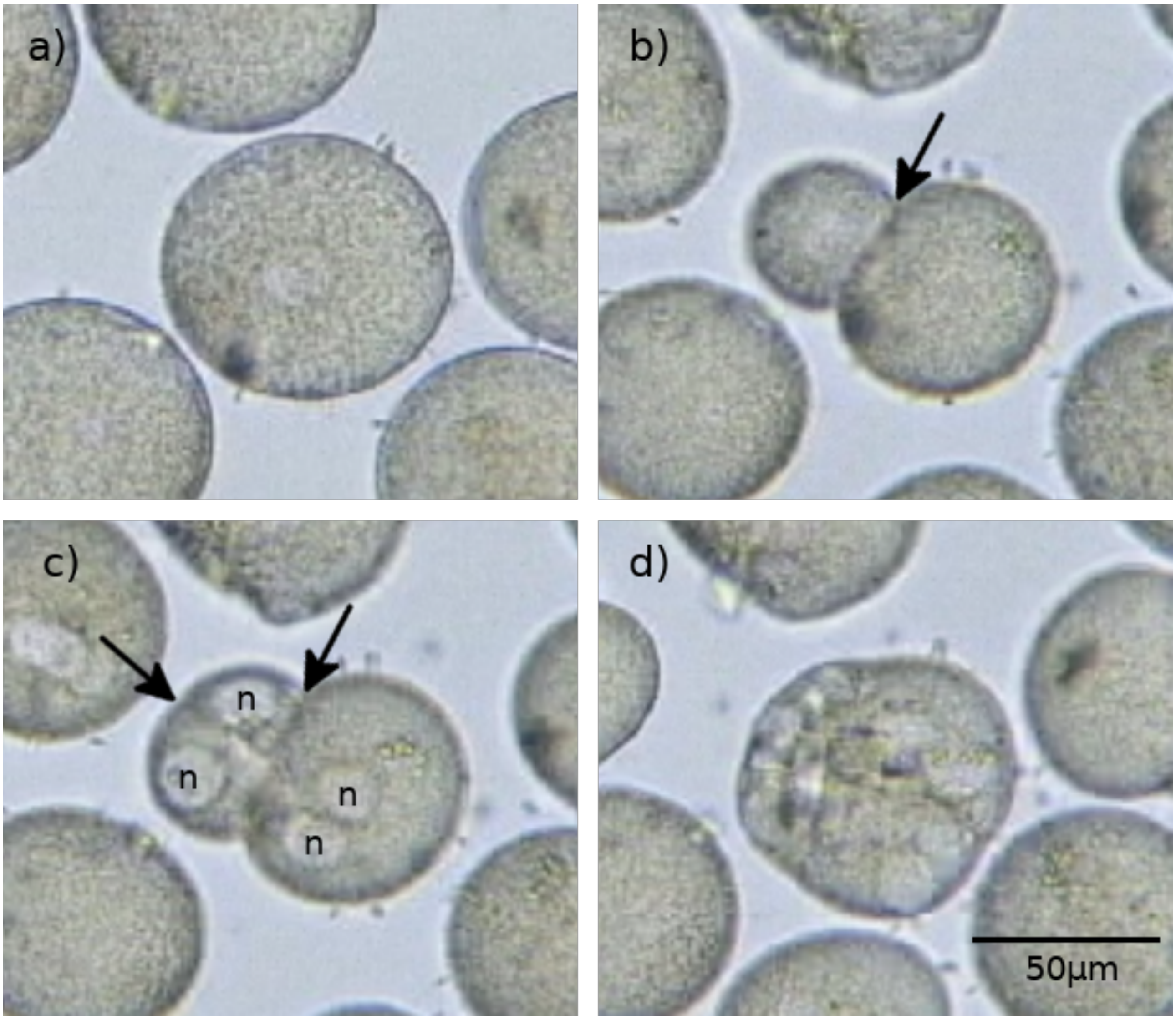

**Figure S32 -** Development under high salt conditions. a) Pre-cleavage zygote. b) Two-cell embryo approaching second division with no cleavage cavities apparent. c) Four-cell embryo with prominent nuclei (n) but lacking cleavage cavities. d) Eight-cell embryo also lacking cleavage cavities. Arrows indicate cleavage planes. Nuclei are marked with an n. Cleavage planes are less clear in the eight-cell embryo because or cell orientation and image resolution.

Zygotes reduced in volume prior to the first cleavage, most likely as a result of osmotic water loss across the fertilisation envelope and plasma membrane (Fig. S33).

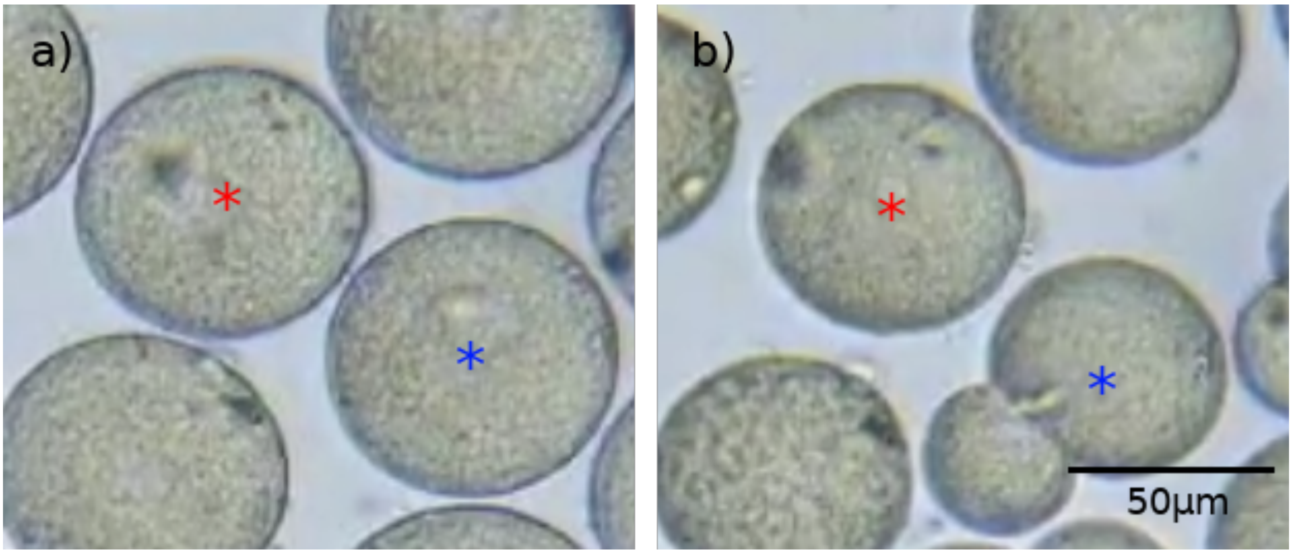

**Figure S33 -** Decrease in volume of unfertilised eggs over time. a) Shortly after fertilisation both the zygote (blue star) and the unfertilised egg (red star) are equal in volume. b) Shortly after the first cleavage of the zygote, the diameter and volume of the unfertilised egg has visibly reduced.

**SM 9.3 - Development and cleavage cavity formation under hypoosmotic conditions**

To test how cleavage cavity formation was affected by low osmotic conditions, embryos were raised in FRW diluted in reverse-osmosis (RO) water. Eggs were collected and processed as per SM 8.2 before being transferred to hypoosomotic media. When subjected to pure RO water, most zygotes failed to cleave. Those that did cleave experienced dissociation of the resulting daughter cells and eventually lysis approximately two hours after fertilisation. Embryos treated with 25% FRW in RO water developed normally and produced large cleavage cavities similar to those in ambient conditions (Fig. S34).

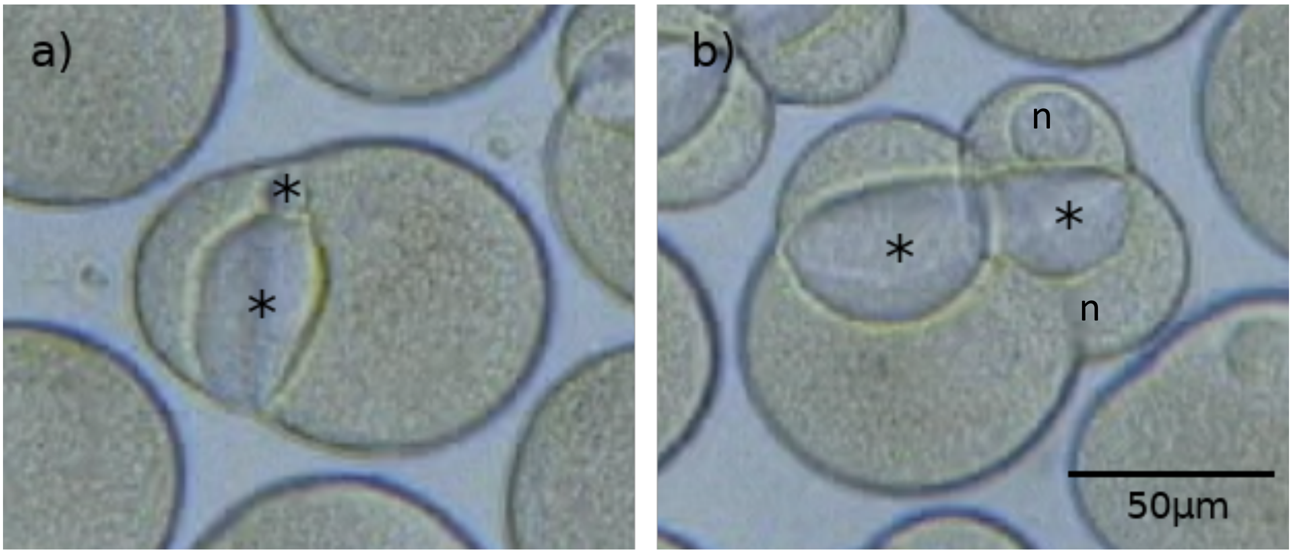

**Figure S34 -** Development under low salt conditions. a) Two-cell embryo with one large and one small cleavage cavity. b) Four-cell embryo with two large cleavage cavities located between the newly divided cells. Cleavage cavities are marked with a *. Nuclei are marked with an n.

### References

1. G. Rosenberg, A nomenclatural review of *Dreissena* (Bivalvia: Dreissenidae), with identification of the quagga mussel as *Dreissena bugensis*. **51**, 1474–1484 (1994).
2. T. W. Therriault, M. F. Docker, M. I. Orlova, D. D. Heath, H. J. MacIsaac, Molecular resolution of the family Dreissenidae (Mollusca: Bivalvia) with emphasis on Ponto-Caspian species, including first report of *Mytilopsis leucophaeata* in the Black Sea basin. *Mol. Phylogenet. Evol.* **30**, 479–489 (2004).
3. D. Micu, A. Telembici, First record of *Dreissena bugensis* (Andrusov, 1897) from the Romanian stretch of River Danube. *Abstr. Int. Symp. Malacol. August 19–22, 2004. Sibiu, Rom.* (2004).
4. M. Rakovic, N. Popovic, V. Kalafatic, V. Martinovic-Vitanovic, Spreading of *Dreissena rostriformis bugensis* (Andrusov, 1897) in the Danube River (Serbia). **65**, 349–357 (2013).
5. G. van der Velde, D. Platvoet, Quagga mussels *Dreissena rostriformis bugensis* (Andrusov, 1897) in the Main River (Germany). *Aquat. Invasions*. **2**, 261–264 (2007).
6. V. L. Gonzalez *et al.*, A phylogenetic backbone for Bivalvia: an RNA-seq approach. *Proc. R. Soc. B Biol. Sci.* **282**, 20142332–20142332 (2015).
7. I. Ebersberger, S. Strauss, A. Von Haeseler, HaMStR: Profile hidden markov model based search for orthologs in ESTs. *BMC Evol. Biol.* **9** (2009).
8. K. Katoh, D. M. Standley, MAFFT multiple sequence alignment software version 7: Improvements in performance and usability. *Mol. Biol. Evol.* **30**, 772–780 (2013).
9. S. R. Eddy, Accelerated profile HMM searches. *PLoS Comput. Biol.* **7** pge.1002195 (2011).
10. P. Kück, K. Meusemann, FASconCAT: Convenient handling of data matrices. *Mol. Phylogenet. Evol.* **56**, 1115–1118 (2010).
11. M. N. Price, P. S. Dehal, A. P. Arkin, FastTree 2 - Approximately maximum-likelihood trees for large alignments. *PLoS One*. **5** pg. e9490 (2010).
12. J. Sambrook, *Molecular cloning : a laboratory manual.* (Cold Spring Harbor Laboratory, Cold Spring Harbor, N.Y, 2001).
13. A. M. Bolger, M. Lohse, B. Usadel, Trimmomatic: a flexible trimmer for Illumina sequence data. *Bioinformatics*. **30**, 2114–2120 (2014).
14. G. Marçais, C. Kingsford, A fast, lock-free approach for efficient parallel counting of occurrences of k-mers. *Bioinformatics*. **27**, 764–70 (2011).
15. G. W. Vurture *et al.*, GenomeScope: Fast reference-free genome profiling from short reads. *Bioinformatics*. **33**, 2202–2204 (2017).

16. T. R. Gregory, Genome size estimates for two important freshwater molluscs, the zebra mussel (*Dreissena polymorpha*) and the schistosomiasis vector snail (*Biomphalaria glabrata*). *Genome*. **46**, 841–4 (2003).

17. B. Liu *et al.*, *arXiv*, in press (available at http://arxiv.org/abs/1308.2012).

18. R. Kajitani *et al.*, Efficient de novo assembly of highly heterozygous genomes from whole-genome shotgun short reads. *Genome Res.* **24**, 1384–95 (2014).

19. L. P. Pryszcz, T. Gabaldón, Redundans : an assembly pipeline for highly heterozygous genomes. *Nucleic Acids Res.* **44**, e113 (2016).

20. S. Wang *et al.*, Scallop genome provides insights into evolution of bilaterian karyotype and development. *Nat. Ecol. Evol.* **1**, 0120 (2017).

21. B. Langmead, S. L. Salzberg, Fast gapped-read alignment with Bowtie 2. *Nat. Methods*. **9**, 357–359 (2012).

22. D. R. Laetsch, M. L. Blaxter, BlobTools: Interrogation of genome assemblies. *F1000Research*. **6**, 1287 (2017).

23. A. F. A. Smit, R. Hubley, P. Green, *RepeatMasker Open-4.0*. 2013-2015 <http://www.repeatmasker.org>.

24. J. Liu *et al.*, BinPacker: Packing-Based De Novo Transcriptome Assembly from RNA-seq Data. *PLoS Comput. Biol.* **12**, 1–15 (2016).

25. D. R. Zerbino, E. Birney, Velvet: algorithms for de novo short read assembly using de Bruijn graphs. *Genome Res.* **18**, 821–9 (2008).

26. B. Bushnell, *BBMap.* (2017) <https://sourceforge.net/projects/bbmap/>.

27. B. J. Haas *et al.*, De novo transcript sequence reconstruction from RNA-seq using the Trinity platform for reference generation and analysis. *Nat. Protoc.* **8**, 1494–1512 (2013).

28. R. C. Edgar, Search and clustering orders of magnitude faster than BLAST. *Bioinformatics*. **26**, 2460–1 (2010).

29. T. D. Wu, C. K. Watanabe, GMAP: A genomic mapping and alignment program for mRNA and EST sequences. *Bioinformatics*. **21**, 1859–1875 (2005).

30. A. Dobin *et al.*, STAR: Ultrafast universal RNA-seq aligner. *Bioinformatics*. **29**, 15–21 (2013).

31. M. Pertea *et al.*, StringTie enables improved reconstruction of a transcriptome from RNA-seq reads. *Nat. Biotechnol.* **33**, 290–5 (2015).

32. M. Stanke, B. Morgenstern, AUGUSTUS: A web server for gene prediction in eukaryotes that allows user-defined constraints. *Nucleic Acids Res.* **33**, 465–467 (2005).

33. I. Korf, Gene finding in novel genomes. *BMC Bioinformatics*. **5**, 59 (2004).

34. E. Birney, M. Clamp, R. Durbin, GeneWise and Genomewise. *Genome Res.* **14**, 988–995 (2004).

35. B. J. Haas *et al.*, Automated eukaryotic gene structure annotation using EVidenceModeler and the Program to Assemble Spliced Alignments. *Genome Biol.* **9**, R7 (2008).

36. N. L. Bray, H. Pimentel, P. Melsted, L. Pachter, Near-optimal probabilistic RNA-seq quantification. *Nat. Biotechnol.* **34**, 525–527 (2016).

37. F. A. Simão, R. M. Waterhouse, P. Ioannidis, E. V Kriventseva, E. M. Zdobnov, BUSCO: assessing genome assembly and annotation completeness with single-copy orthologs. *Bioinformatics*. **31**, 3210–3212 (2015).

38. M. Kanehisa, Y. Sato, M. Kawashima, M. Furumichi, M. Tanabe, KEGG as a reference resource for gene and protein annotation. *Nucleic Acids Res.* **44**, D457–D462 (2016).

39 Y. Moriya, M. Itoh, S. Okuda, A. C. Yoshizawa, M. Kanehisa, KAAS: An automatic genome annotation and pathway reconstruction server. *Nucleic Acids Res.* **35**, 182–185 (2007).

40. R-Core-Team, R: A Language and Environment for Statistical Computing. *Vienna, Austria R Found. Stat. Comput.* **1** (2015).

41. P. J. Kersey *et al.*, Ensembl Genomes 2018: An integrated omics infrastructure for non-vertebrate species. *Nucleic Acids Res.* **46**, D802–D808 (2018).

42. O. Simakov *et al.*, Insights into bilaterian evolution from three spiralian genomes. *Nature*. **493**, 526–531 (2013).

43. J. Sun *et al.*, Data from: Adaptation to deep-sea chemosynthetic environments as revealed by mussel genomes. *Nat. Ecol. Evol.* (2017).

44. G. Zhang *et al.*, The oyster genome reveals stress adaptation and complexity of shell formation. *Nature*. **490**, 49–54 (2012).

45. M. Uliano-Silva *et al.*, A hybrid-hierarchical genome assembly strategy to sequence the invasive golden mussel *Limnoperna fortunei*. *Gigascience*. **7**, gix128 (2018).

46. T. Takeuchi *et al.*, Draft genome of the pearl oyster *Pinctada fucata*: a platform for understanding bivalve biology. *DNA Res.* **19**, 117–30 (2012).

47. N. H. Putnam *et al.*, The amphioxus genome and the evolution of the chordate karyotype. *Nature*. **453**, 1064–1071 (2008).

48. C. B. Albertin *et al.*, The octopus genome and the evolution of cephalopod neural and morphological novelties. *Nature*. **524**, 220–224 (2015).

49. R. G. S. P. Consortium *et al.*, Genome sequence of the Brown Norway rat yields insights into mammalian evolution. *Nature*. **428**, 493 (2004).

50. N. H. Putnam *et al.*, Sea anemone genome reveals ancestral eumetazoan gene repertoire and genomic organization. *Science*. **317**, 86–94 (2007).

51. C. M. Adema *et al.*, Whole genome analysis of a schistosomiasis-transmitting freshwater snail. *Nat. Commun.* **8**, 15451 (2017).

52. E. Sodergren *et al.*, *The genome of the sea urchin* Strongylocentrotus purpuratus. *Science.* **314**, 941–952 (2006).

53. M. D. Adams *et al.*, *The genome sequence of* Drosophila melanogaster. Science. **287**, 2185–2195 (2000).

54. W. Li, A. Godzik, Cd-hit: A fast program for clustering and comparing large sets of protein or nucleotide sequences. *Bioinformatics*. **22**, 1658–1659 (2006).

55. Y. Peng *et al.*, IDBA-tran: a more robust de novo de Bruijn graph assembler for transcriptomes with uneven expression levels. *Bioinformatics*. **29**, i326-34 (2013).

56. A. Larsson, AliView: A fast and lightweight alignment viewer and editor for large datasets. *Bioinformatics*. **30**, 3276–3278 (2014).

57. A. Criscuolo, S. Gribaldo, BMGE (Block Mapping and Gathering with Entropy): A new software for selection of phylogenetic informative regions from multiple sequence alignments. *BMC Evol. Biol.* **10** (2010).

58. R. N. Finn, J. Cerdà, Evolution and Functional Diversity of Aquaporins. *Biol. Bull.* **229**, 6–23 (2015).

59. R. N. Finn, F. Chauvigné, J. A. Stavang, X. Belles, J. Cerdà, Insect glycerol transporters evolved by functional co-option and gene replacement. *Nat. Commun.* **6**, 1–7 (2015).

60. E. Kosicka, D. Grobys, H. Kmita, A. Lesicki, J. R. Pieńkowska, Putative new groups of invertebrate water channels based on the snail *Helix pomatia L.* (Helicidae) MIP protein identification and phylogenetic analysis. *Eur. J. Cell Biol.* **95**, 543–551 (2016).

61. N. Kaufmann *et al.*, Developmental expression and biophysical characterization of a *Drosophila melanogaster* aquaporin. *Am. J. Physiol. Physiol.* **289**, C397–C407 (2005).

62. K.-S. Lee *et al.*, Molecular cloning and expression of a cDNA encoding the aquaporin homologue from the firefly, *Pyrocoelia rufa*. *Korean J. Entomol*. **31**, 269–279 (2001).

63. N. Kataoka, S. Miyake, M. Azuma, Molecular characterization of aquaporin and aquaglyceroporin in the alimentary canal of *Grapholita molesta* (the oriental fruit moth) - comparison with *Bombyx mori* aquaporins. *J. Insect Biotechnol. Sericology*. **78**, 81–90 (2009).

64. N. Kataoka, S. Miyake, M. Azuma, Aquaporin and aquaglyceroporin in silkworms, differently expressed in the hindgut and midgut of *Bombyx mori*. *Insect Mol. Biol.* **18**, 303–314 (2009).

65. I. S. Wallace *et al.*, *Acyrthosiphon pisum* AQP2: A multifunctional insect aquaglyceroporin. *Biochim. Biophys. Acta - Biomembr.* **1818**, 627–635 (2012).

66. L. L. Drake, S. D. Rodriguez, I. A. Hansen, Functional characterization of aquaporins and aquaglyceroporins of the yellow fever mosquito, *Aedes aegypti*. *Sci. Rep.* **5**, 1–7 (2015).

67. A. Madeira *et al.*, Human Aquaporin-11 is a water and glycerol channel and localizes in the vicinity of lipid droplets in human adipocytes. *Obesity*. **22**, 2010–2017 (2014).

68. D. J. Colgan, R. P. Santos, A phylogenetic classification of gastropod aquaporins. *Mar. Genomics* **38**, 59-65 (2017).

69. M. Toei, R. Saum, M. Forgac, Regulation and isoform function of the V-ATPases. *Biochemistry*. **49**, 4715–4723 (2010).

70. M. E. Maxson, S. Grinstein, The vacuolar-type H+-ATPase at a glance - more than a proton pump. *J. Cell Sci.* **127**, 4987–4993 (2014).

71. S. Kawasaki-Nishi, K. Bowers, T. Nishi, M. Forgac, T. H. Stevens, The amino-terminal domain of the vacuolar proton-translocating ATPase a subunit controls targeting and in vivo dissociation, and the carboxyl-terminal domain affects coupling of proton transport and ATP hydrolysis. *J. Biol. Chem.* **276**, 47411–47420 (2001).

72. M. Forgac, Vacuolar ATPases: Rotary proton pumps in physiology and pathophysiology. *Nat. Rev. Mol. Cell Biol.* **8**, 917–929 (2007).

73. A. N. Smith *et al.*, Mutations in ATP6N1B, encoding a new kidney vacuolar proton pump 116-kD subunit, cause recessive distal renal tubular acidosis with preserved hearing. *Nat. Genet.* **26**, 71–75 (2000).

74. A. N. Smith *et al.*, Molecular Cloning and Characterization of Atp6n1b. A novel fourth murine vacuolar H+-ATPase a-subunit gene. *J. Biol. Chem.* **276**, 42382–42388 (2001).

75. C. L. Brett, M. Donowitz, R. Rao, Evolutionary origins of eukaryotic sodium / proton exchangers. *Am. J. Physiol. - Cell Physiol.* **288**, C223–C239 (2005).

76. A. Waterhouse *et al.*, SWISS-MODEL: Homology modelling of protein structures and complexes. *Nucleic Acids Res.* **46**, W296–W303 (2018).

77. J. Yu, A. J. Yool, K. Schulten, E. Tajkhorshid, Mechanism of Gating and Ion Conductivity of a Possible Tetrameric Pore in Aquaporin-1. *Structure*. **14**, 1411–1423 (2006).

78. D. Alberga *et al.*, A new gating site in human aquaporin-4: Insights from molecular dynamics simulations. *Biochim. Biophys. Acta - Biomembr.* **1838**, 3052–3060 (2014).

79. M. Kourghi, M. L. De Ieso, S. Nourmohammadi, J. V. Pei, A. J. Yool, Identification of loop D domain amino acids in the human aquaporin-1 channel involved in activation of the ionic conductance and inhibition by AqB011. *Front. Chem.* **6**, 1–12 (2018).

80. C. Rodrigues *et al.*, Rat aquaporin-5 is pH-gated induced by phosphorylation and is implicated in oxidative stress. *Int. J. Mol. Sci.* **17** (2016).

81. S. Costantini, G. Colonna, A. M. Facchiano, ESBRI: A web server for evaluating salt bridges in proteins. *Bioinformation*. **3**, 137–138 (2008).

82. D. E. Anderson, W. J. Becktel, F. W. Dahlquist, pH-Induced denaturation of proteins: A single salt bridge contributes 3-5 kcal/mol to the free energy of folding of T4 lysozyme. *Biochemistry*. **29**, 2403–2408 (1990).

83. N. V. Di Russo, D. A. Estrin, M. A. Martí, A. E. Roitberg, pH-dependent conformational changes in proteins and their effect on experimental pKas: The case of nitrophorin 4. *PLoS Comput. Biol.* **8** (2012).

84. T. J. Wheeler, J. Clements, R. D. Finn, Skylign: A tool for creating informative, interactive logos representing sequence alignments and profile hidden Markov models. *BMC Bioinformatics*. **15**, 1–9 (2014).

85. V. D. Vacquier, B. Brandriff, C. G. Glabe, The effect of soluble egg jelly on the fertilizability of acid-dejellied sea urchin eggs. *Develop. Growth Differ*. **21**, 47–60 (1979).
